# Frontal Cortical Hemodynamic Responses and Inter-Brain Synchrony During Naturalistic Co-Viewing of Differing Movie Types

**DOI:** 10.64898/2026.11.17.767060

**Authors:** Xusheng Fang, Mark Byrd, Khurshid Ahmad, Daniela Tropea, Shane M. O’Mara

## Abstract

Simultaneous recording during shared stimulus viewing by dyads may reveal underlying cortical hemodynamic alignment arising from common stimulus processing, co-presence of others, and coordinated or correlated movement. Here, we conducted frontal cortical functional near-infrared spectroscopy (fNIRS) recordings from 46 dyads (23 friend and 23 stranger pairs; 92 female participants) during viewing of *audio, comic*, and *horror* movie stimuli (fifty mins duration). We measured non-partner inter-subject correlations (ISC) and compared wavelet transform coherence (WTC) in actual and stimulus-matched pseudo-dyads. Primary oxygenated hemoglobin (HbO) ISC was positive in all three *audio, comic*, and *horror* conditions. Actual dyads had higher HbO WTC during *comic*, including after removal of 60 seconds from each boundary. This effect did not survive false discovery rate (FDR) correction after individual head-motion regression in 44 dyads. Complementary deoxygenated hemoglobin (HbR) analyses showed positive ISC and a *comic* real-minus-pseudo-WTC advantage that persisted after motion regression. Neither chromophore showed a consistent familiarity effect. Overall, our findings indicate shared hemodynamic synchrony across participants and a small but significant increase in inter-brain synchrony in actual dyads compared with pseudo-dyads during the *comic* condition, observed in both HbO and HbR signals. Although the HbR effect persisted after adjustment for measured head motion, potential contributions from shared-session factors and systemic physiological influences cannot be ruled out, highlighting the need for caution when interpreting inter-brain synchrony during naturalistic co-viewing.

## 1 Introduction

While many social experiences involve two people responding to the same events, social cognition is often studied in individuals observing the actions of others (Amodio and Frith, 2006; Schilbach et al., 2013; Redcay and Schilbach, 2019). Two people can respond similarly to a common stimulus without interacting with each other (e.g., in classrooms) or other social settings (e.g. in movie theatres) where there may be shared sensory inputs (Dikker et al., 2017; Pérez et al., 2017; Piazza et al., 2020). The same film or narrative can therefore produce correlated activity across independently recorded participants (measured by inter-subject correlations or ISC; Hasson et al., 2004; Nummenmaa et al., 2012; Nastase et al., 2019; Simony et al., 2016). Seeming inter-brain coherence may therefore reflect common stimulus processing rather than, or in addition to, social or other coordination between partners (Dumas et al., 2010; Hasson et al., 2012).

Pseudo-dyad comparisons can help address this ambiguity; synchrony can arise when recordings from people who did not interact are paired analytically (Burgess, 2013), and shared task structure is a recognized confound in hyperscanning research (Hamilton, 2021). Comparing actual partners with stimulus-matched pseudo-dyads tests whether coherence is greater between people recorded together. But such a difference cannot establish direct neural influence: actual partners also share a physical setting and the same stimulus session, whereas pseudo-dyads do not (Leahy et al., 2025). Relationship familiarity may also affect shared responses. Friends can show similar neural activity when watching the same material separately (Parkinson et al., 2018). Other studies have related friendship or close relationships to neural responses and coordination during social tasks (Güroğlu et al., 2008; Meyer et al., 2013; Kinreich et al., 2017; Pan et al., 2017; Reindl et al., 2018). It remains unclear whether friends show additional coherence when watching the same stimulus together, beyond similarity of stimulus processing associated with the common stimulus.

Hyperscanning involves recording brain activity from multiple people simultaneously engaged in the same task (Montague et al., 2002; Czeszumski et al., 2020), offering the possibility of investigating neural activity in social settings. fNIRS (functional near-infrared spectroscopy) is a non-invasive optical method for measuring and imaging changes in relative oxygen cerebral hemoglobin concentration, and thereby measure cerebral hemodynamics (HbR is deoxygenated hemoglobin and HbO is oxygenated hemoglobin). fNIRS capitalizes on the fact that HbO and HbR have distinct near-infrared absorption spectra, allowing measurement of their concentration changes separately, and thereby allowing inferences regarding neural activity through the ongoing dynamics of hemodynamic response. Recent advances in portable fNIRS makes hyperscanning of processes supporting cognition possible in naturalistic settings (Cui et al., 2012; Hirsch et al., 2017; Jiang et al., 2012; Pinti et al., 2020).

Here, using fNIRS, we recorded friend and stranger dyads during naturalistic co-viewing of differing short movie types. Our main questions concerned whether non-partners showed shared hemodynamic responses, if actual (real-life) partners showed greater coherence than stimulus-matched pseudo-dyads, and if coherence differed with familiarity. Phase and sensitivity analyses examined alternative explanations, including stimulus boundaries and head motion. Complementary HbR analyses assessed whether the findings were restricted to HbO.

## 2 Materials and Methods

### 2.1 Participants

Ninety-two healthy female participants formed 46 dyads (mean age = 21.82 years, SD = 3.76, range = 18-38 years). Twenty-three dyads were friends who had known one another for at least three months; the other 23 were strangers with no prior contact. Participants were neurotypical, right-handed, fluent in English, and had normal or corrected-to-normal vision and hearing. They received course credit or a €15 voucher. All provided written informed consent, and the Ethics Committee of the School of Psychology, Trinity College Dublin, approved the study (REAMs reference no. 5291; approval letter dated 19 January 2026).

### 2.2 Experimental Design and Procedure

Participants sat together in front of a shared display for 50 minutes of continuous fNIRS recording. The session included an initial eyes-closed reference period, three naturalistic stimulus blocks, two intervening opportunities for unrestricted conversation, and a final eyes-closed reference period (Figure 1).

**Figure 1.**
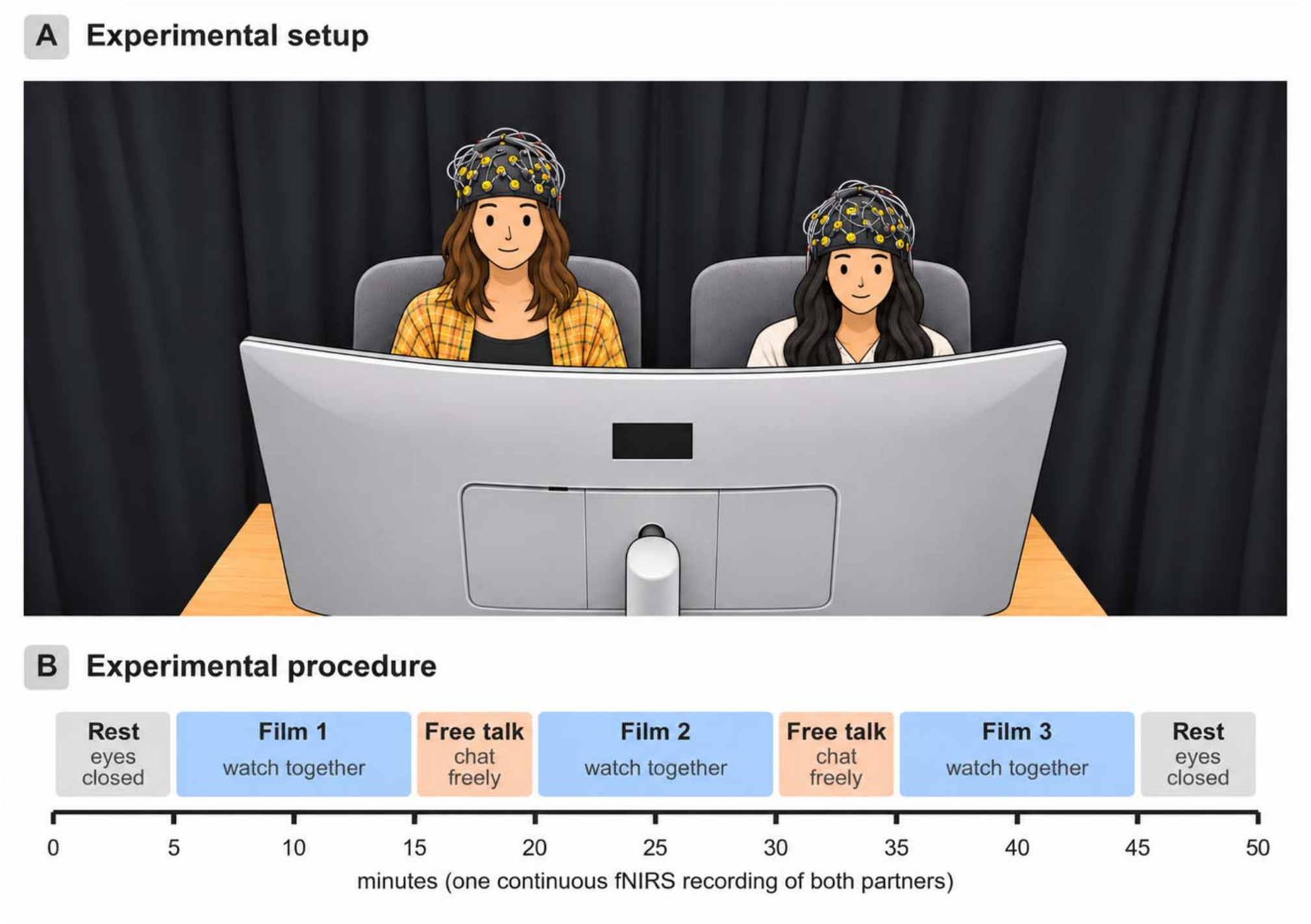
Experimental setup and timeline. (A) Participants sat side by side facing a shared screen while wearing fNIRS caps. (B) The approximately 50-minute recording included two five-minute rest periods, three ten-minute stimulus blocks in counterbalanced order, and two five-minute free-conversation periods.

The recording sequence comprised the following blocks:

- *Initial reference period:* Five minutes with eyes closed.
- *Stimulus blocks:* Three 10-minute conditions (*audio, comic, horror*), presented across dyads, using all six possible orders.
- *Free interaction:* A five-minute eyes-open period between successive stimulus blocks, during which participants could converse freely.
- *Final reference period:* Five minutes with eyes closed. Reference and free-interaction recordings were not included in the stimulus-specific analyses.

Stimulus onset and offset were identified from alternating high (H) and low (L) event markers in each SNIRF file, using the recorded presentation-order code to identify each condition. Partner recordings were aligned within sessions. Each analyzed segment contained 14,975 samples (approximately 600 seconds at 25 Hz).

### 2.3 Data Acquisition and Preprocessing

Frontal hemodynamic signals were recorded simultaneously from both participants using two fNIRS systems (Brite24; Artinis Medical Systems B.V., Elst, the Netherlands), with one 24-channel cap per participant. Each system comprised 10 light sources nominally operating at wavelengths of 760 and 850 nm (with calibrated wavelengths 763.8/844.9 and 756.0/842.8 nm for the two devices) and eight detectors, arranged in a frontal montage with a nominal source-detector separation of 30 mm. Optical intensity was sampled at approximately 25 Hz. Using MNE-based Python routines, we converted intensity to optical density and calculated the scalp coupling index (SCI) before motion correction. Channels were retained when the lower SCI across wavelengths was at least .70. We applied temporal derivative distribution repair (TDDR; Fishburn et al., 2019) and converted optical density to oxygenated (HbO) and deoxygenated hemoglobin (HbR) using the modified Beer– Lambert law (differential pathlength factor = 6). A zero-phase Hamming-window FIR filter (0.02– 0.10 Hz; transition bandwidths .01 and .02 Hz) was applied to continuous data before extracting stimulus blocks. HbO was the primary measure; HbR was analyzed subsequently as a complementary, exploratory measure. Without short-separation channels or cardiovascular recordings, cortical and systemic components could not be separated fully (Yücel et al., 2021).

### 2.4 Inter-Brain Synchrony and Auxiliary Measurements

Wavelet transform coherence (WTC) was computed over 0.02-0.10 Hz using 64 complex Morlet scales (Grinsted et al., 2004), excluding coefficients inside the cone of influence. The main dyad score averaged channels and overlapping 50-second windows (one-second steps); each window contributed atanh(sqrt(mean valid squared coherence)). We refer to this transformed, phase-insensitive measure as absolute WTC. It is not raw coherence or a baseline-subtracted score. Non-partner ISC was calculated separately from Fisher-z-transformed Pearson correlations; the two metrics are not numerically comparable. Accelerometer and gyroscope data were available for both participants in 44 of the 46 dyads. One participant in each of the remaining two dyads had unusable inertial-sensor data. Motion sensitivity analyses were therefore restricted to the 44 dyads with complete recordings from both participants. (Supplementary Methods).

### 2.5 Statistical Analysis

We assessed (Q1) non-partner ISC, (Q2) actual-versus-pseudo-dyad WTC, (Q3) friend-versus-stranger WTC, and (Q4) phase organization. For ISC, actual partners were excluded from pairing; each participant’s mean ISC across eligible non-partners was tested against zero using a two-sided Wilcoxon signed-rank test (n = 92, non-independent), with an additional sign-flip test that accounts for non-independence of samples (Chen et al., 2016).

For WTC, the 46 actual A-B dyads were compared with all 2,070 possible non-partner A-B combinations. One-sided significance was assessed using 10,000 random one-to-one re-matchings of the 46 A and 46 B participants, excluding their actual partners. Pseudo-pair means were descriptive; the 2,070 combinations were not treated as independent observations. The primary re-matching was stimulus-matched but not presentation-order-matched. Friendship comparisons used one WTC value per actual dyad and a two-sided Mann–Whitney U test (23 versus 23). Phase angles were examined with Rayleigh and V-tests; exploratory channel-, time-, and phase-specific analyses are described with their correction families in the Supplementary Results.

Benjamini–Hochberg false discovery rate (FDR) correction was applied across the three stimulus conditions within each condition-level analysis and separately within the reported exploratory families (Benjamini and Hochberg, 1995). Pooled results across conditions were descriptive. Motion sensitivity analyses compared unchanged HbO, HbO after own-motion regression, and coherence adjusted for between-partner movement coupling in the 44-dyad sensor subset. Analogous HbR motion checks were exploratory (Supplementary Methods).

Two-sided 95% percentile confidence intervals (CIs) were computed from 20,000 stratified resamples of intact dyads, preserving the friend-stranger balance. The real-versus-pseudo contrast was re-estimated in each resample without treating overlapping pseudo-pairs as independent. These CIs describe sampling uncertainty and are not inversions of the original re-matching or rank-based tests; the CIs were not multiplicity-adjusted. Circular phase intervals used analogous dyad resampling (Supplementary Methods).

The complementary HbR analyses used the same Q1-Q4 definitions, dyad-level uncertainty estimation, and respective permutation or rank-based tests. Each three-stimulus HbR family was FDR-corrected separately; a six-test HbO/HbR correction was also examined for the actual-versus-pseudo-WTC comparisons. HbR WTC and phase computations involved source-calibrated lower-sampling-rate wavelet estimates (Supplementary Methods).

### 2.6 Exploratory stimulus-boundary and FIR-input sensitivity analyses

We examined whether the findings depended on stimulus boundaries by repeating ISC and WTC analyses after excluding 30, 60, or 120 seconds from each end of every block. Only WTC windows fully inside the retained interval were used. We also tested alternative signal extensions outside the *comic* block during FIR filtering and, separately, re-matching restricted to the same presentation order. All these checks were exploratory. FDR correction was applied across conditions within each trimming threshold, but not across the different thresholds themselves. Details are given in the Supplementary Methods.

Boundary analyses used HbO sampled at approximately 5 Hz. The reconstructed WTC scores closely agreed with full-rate 25-Hz scores (mean absolute difference = .000032 across 6,348 condition-by-pair estimates; Supplementary Methods).

## 3 Results

All primary HbO comparisons included 46 dyads. Complementary HbR comparisons used the same 46 dyads; motion sensitivity analyses included 44 dyads with usable sensors in both partners.

### Q1 Shared responses across non-partner participants

Non-partner ISC was positive in *audio* (mean Fisher z = .013), *comic* (.031), and *horror* (.044; Table 1 and Figure 2). All three Wilcoxon tests survived FDR correction (adjusted p ≤ 1.0 × 10/□^11^), as did the sign-flip test (p ≤ .0001). The stimulus means were not compared directly, so their ordering is descriptive. Channel-level analyses also showed positive ISC after correction (largest adjusted p = .017).

**Table 1.**
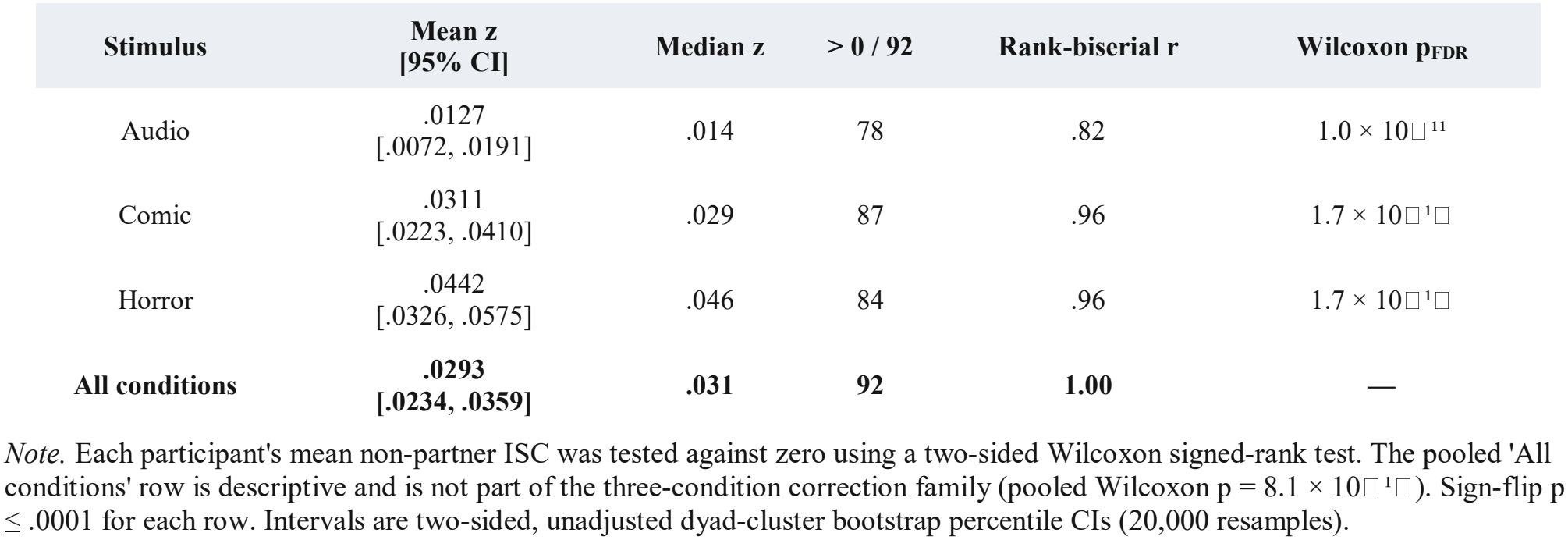
Non-partner HbO inter-subject correlation (ISC) by stimulus.

| **Stimulus** | **Mean z**<br>**[95% CI]** | **Median z** | **> 0 / 92** | **Rank-biserial r** | **Wilcoxon p<sub>FDR</sub>** |
|---|---|---|---|---|---|
| Audio | .0127<br>[.0072, .0191] | .014 | 78 | .82 | 1.0 × 10□<sup>11</sup> |
| Comic | .0311<br>[.0223, .0410] | .029 | 87 | .96 | 1.7 × 10□<sup>1</sup>□ |
| Horror | .0442<br>[.0326, .0575] | .046 | 84 | .96 | 1.7 × 10□<sup>1</sup>□ |
| **All conditions** | **.0293**<br>**[.0234, .0359]** | **.031** | **92** | **1.00** | **—** |

*Note.* Each participant's mean non-partner ISC was tested against zero using a two-sided Wilcoxon signed-rank test. The pooled 'All conditions' row is descriptive and is not part of the three-condition correction family (pooled Wilcoxon p = 8.1 × 10□<sup>1</sup>□). Sign-flip p ≤ .0001 for each row. Intervals are two-sided, unadjusted dyad-cluster bootstrap percentile CIs (20,000 resamples).

**Figure 2.**
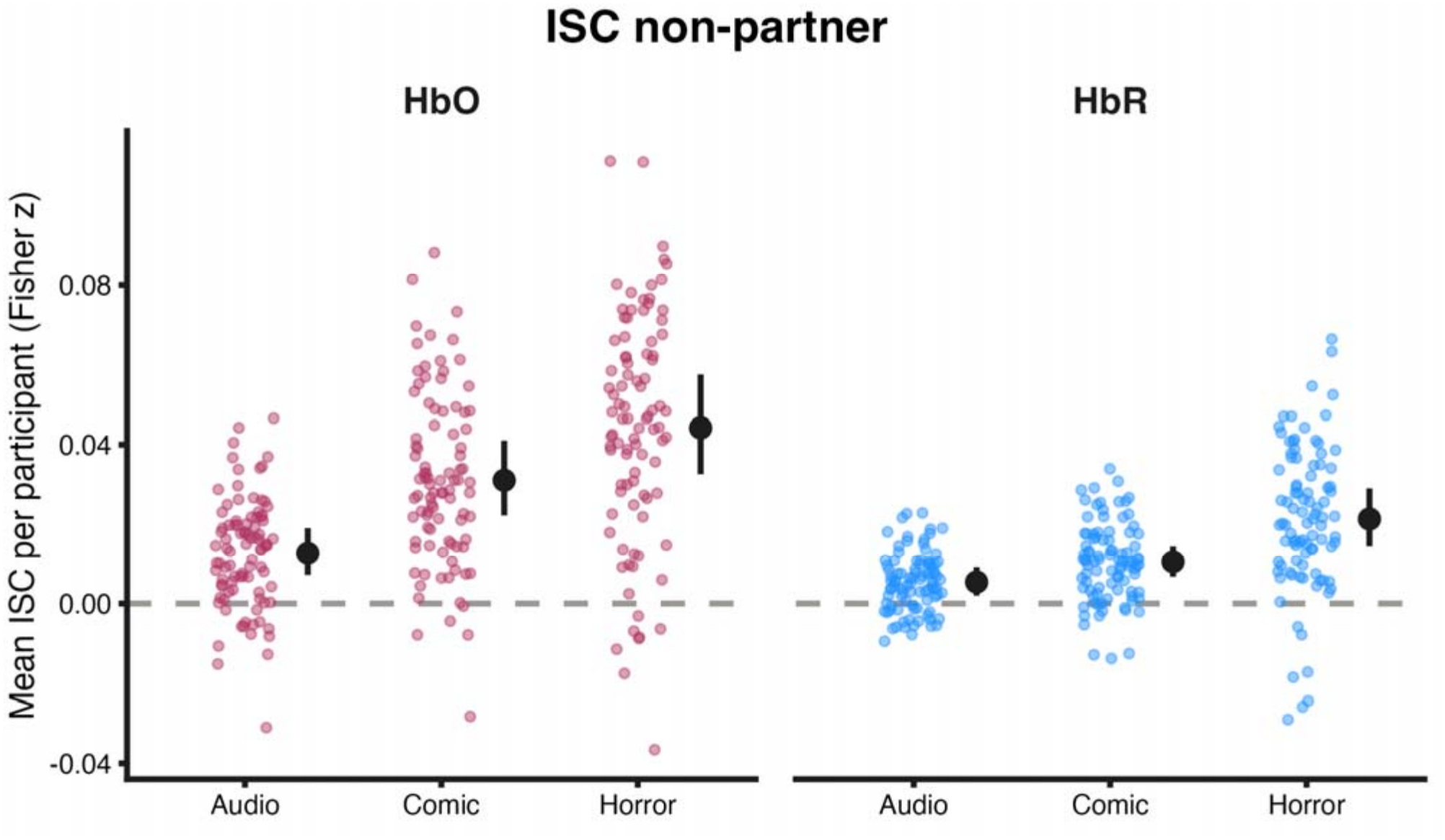
Mean non-partner HbO and HbR ISC (Fisher z) by stimulus (n = 92). Error bars show two-sided 95% dyad-cluster bootstrap CIs. Between-stimulus differences were not tested.

After removing 60 seconds from either end of each block, mean ISC remained positive in all three conditions (z = .01220, .03516, and .04701; all Wilcoxon p ≤ 4.3 × 10/□ ^1^/□; Supplementary Table S1). Thus, the positive ISC did not depend solely on block transitions.

Complementary HbR ISC was also positive in *audio, comic* and *horror* (mean Fisher z = .0054, .0105, and .0214; all FDR-adjusted p < .001; Supplementary Table S9).

### Q2 Real versus stimulus-matched pseudo-dyads

Actual dyads had slightly higher mean WTC than stimulus-matched pseudo-dyads in all three conditions (Table 2; Figure 3). The difference survived FDR correction only in *comic* (real = .7082, pseudo = .7021, difference = +.0062, 95% CI [.0019, .0103], one-sided adjusted p = .036). Neither *audio* (adjusted p = .246) nor *horror* (adjusted p = .083) was significant. The pooled result was descriptive and all differences were sensitive to motion adjustment (Supplementary Table S7).

**Table 2.**
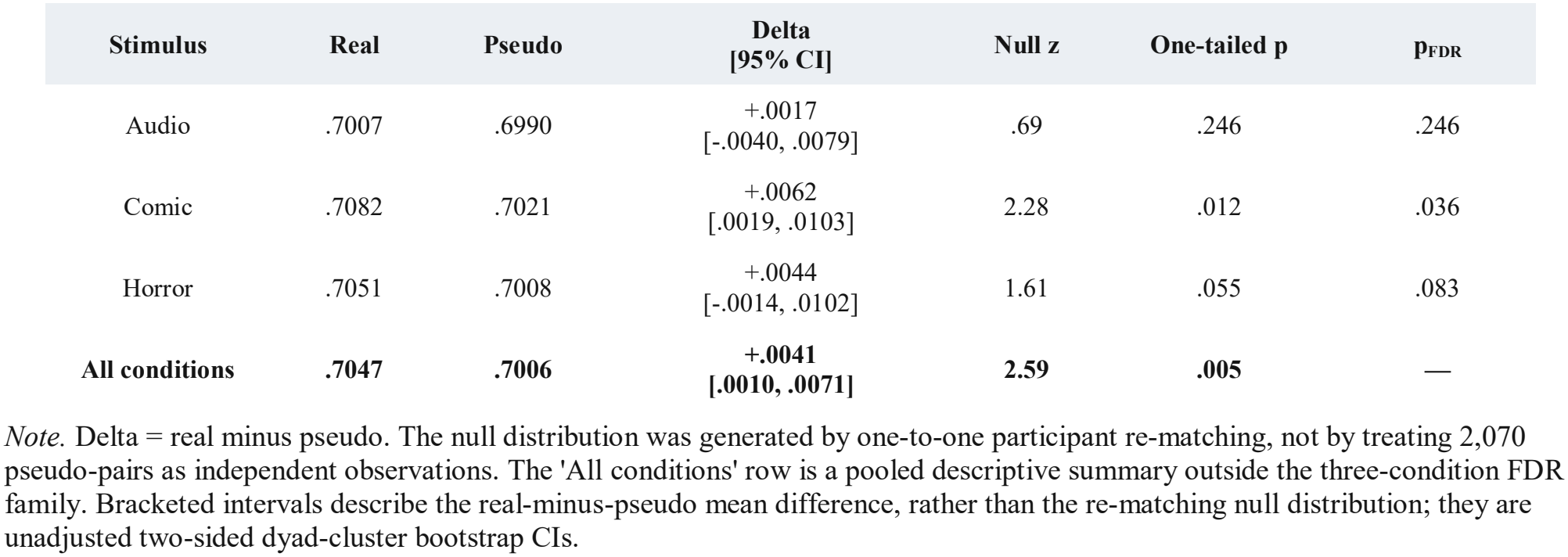
HbO WTC in real dyads versus all 2,070 stimulus-matched pseudo-pairings.

| **Stimulus** | **Real** | **Pseudo** | **Delta**<br>**[95% CI]** | **Null z** | **One-tailed p** | **p<sub>FDR</sub>** |
|---|---|---|---|---|---|---|
| Audio | .7007 | .6990 | +.0017<br>[-.0040, .0079] | .69 | .246 | .246 |
| Comic | .7082 | .7021 | +.0062<br>[.0019, .0103] | 2.28 | .012 | .036 |
| Horror | .7051 | .7008 | +.0044<br>[-.0014, .0102] | 1.61 | .055 | .083 |
| **All conditions** | **.7047** | **.7006** | **+.0041**<br>**[.0010, .0071]** | **2.59** | **.005** | **—** |

*Note.* Delta = real minus pseudo. The null distribution was generated by one-to-one participant re-matching, not by treating 2,070 pseudo-pairs as independent observations. The 'All conditions' row is a pooled descriptive summary outside the three-condition FDR family. Bracketed intervals describe the real-minus-pseudo mean difference, rather than the re-matching null distribution; they are unadjusted two-sided dyad-cluster bootstrap CIs.

**Figure 3.**
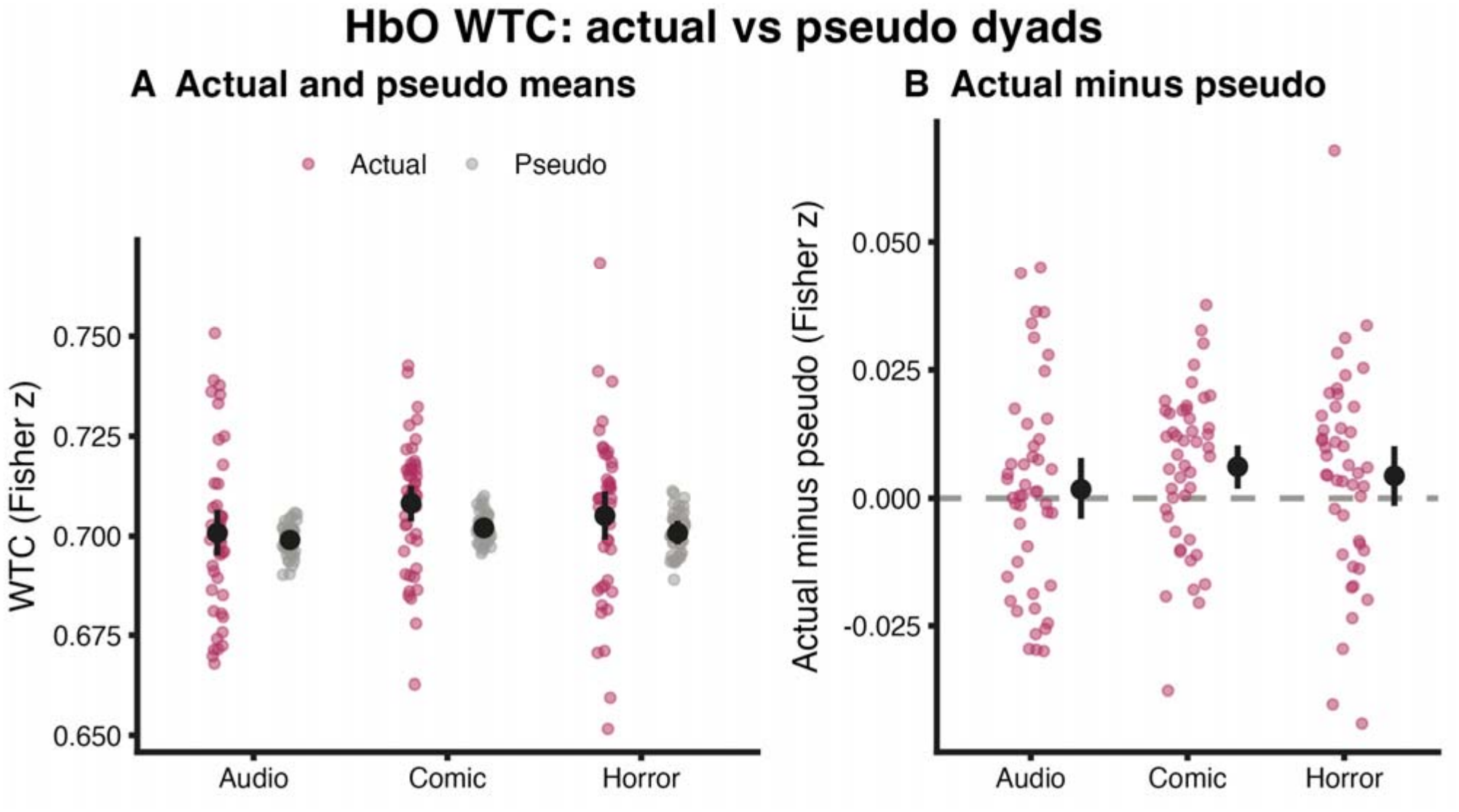
HbO WTC in actual dyads and stimulus-matched pseudo-dyads. (A) Descriptive actual and pseudo means (with 95% dyad-cluster bootstrap CIs). (B) Real-minus-pseudo mean differences with 95% CIs. Pseudo-pair dependence is retained in resampling; inference used participant re-matching and FDR-adjusted p values in Table 2.

The secondary HbR comparison showed a similar *comic*-specific actual-partner advantage (difference = +.0088, 95% CI [.0054, .0123]; FDR-adjusted p = .0003; six-test HbO/HbR adjusted p = .0006). HbR differences in *audio* and *horror* were not significant (Supplementary Table S10 and Figure S4).

### Q2.1 Exploratory sensitivity of real-partner WTC to stimulus boundaries

The *comic* real-minus-pseudo difference remained significant after excluding 30 or 60 seconds from each end of the block (differences = .00653 and .00679; FDR-adjusted p = .0330 and .0366, respectively; Table 3 and Figure 4). At 120 seconds, the estimate was still positive (.00693), but no longer survived FDR correction (p = .0744). These overlapping exploratory checks were not independently corrected across trimming choices.

**Table 3.**
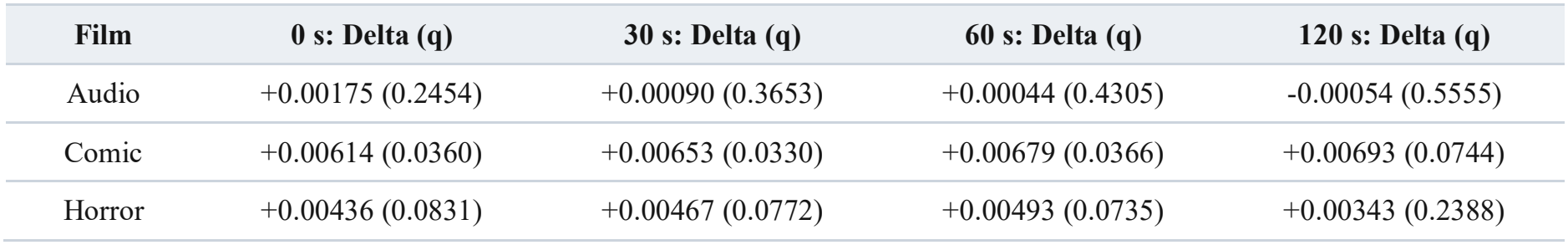
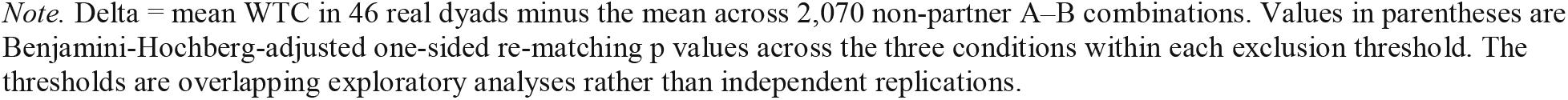
Exploratory real-minus-pseudo HbO WTC after excluding data from both ends of each stimulus block.

| **Film** | **0 s: Delta (q)** | **30 s: Delta (q)** | **60 s: Delta (q)** | **120 s: Delta (q)** |
|---|---|---|---|---|
| Audio | +0.00175 (0.2454) | +0.00090 (0.3653) | +0.00044 (0.4305) | -0.00054 (0.5555) |
| Comic | +0.00614 (0.0360) | +0.00653 (0.0330) | +0.00679 (0.0366) | +0.00693 (0.0744) |
| Horror | +0.00436 (0.0831) | +0.00467 (0.0772) | +0.00493 (0.0735) | +0.00343 (0.2388) |

*Note.* Delta = mean WTC in 46 real dyads minus the mean across 2,070 non-partner A–B combinations. Values in parentheses are Benjamini-Hochberg-adjusted one-sided re-matching p values across the three conditions within each exclusion threshold. The thresholds are overlapping exploratory analyses rather than independent replications.

**Figure 4.**
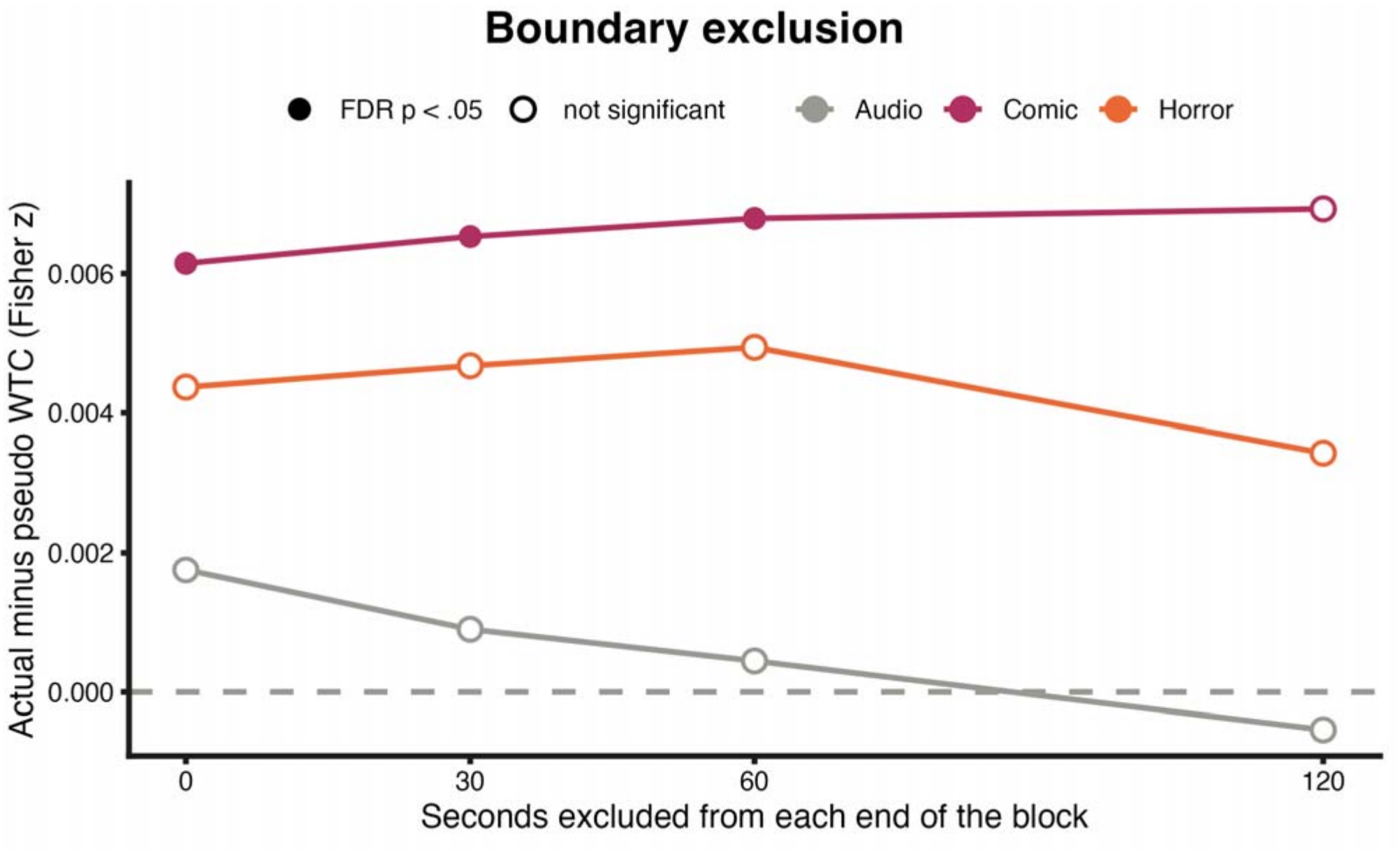
Actual-minus-pseudo HbO WTC differences after excluding data from each stimulus boundary. Each point compares 46 actual dyads with the descriptive pseudo pool. Significance was assessed by one-to-one re-matching; adjusted p values are in Table 3.

For *comic*, replacing the signal outside the block with reflected or constant extensions during FIR filtering produced similar 60-second-trimmed differences (.00682 and .00683 versus .00679 with continuous filtering; Supplementary Table S4 and Figure S2). An order-matched re-matching analysis also gave a positive *comic* difference, but its null distribution differed from the primary comparison (Supplementary Table S3). Neither analysis rules out movement or other factors shared within a session.

### Q3 Social familiarity: friends versus strangers

No friend-versus-stranger WTC difference survived correction (Table 4; Figure 5). Friends showed higher mean coherence in *comic* (difference = +.0094, unadjusted 95% CI [.0004, .0185], uncorrected p = .019), but the FDR-adjusted value was .056. Horror showed a small difference in the opposite direction (−.0034; adjusted p = .742). The pooled comparison was also nonsignificant (p = .180). The more conservative 98.33% Bonferroni CI for the *comic* mean difference [−.0013, .0206] included zero.

**Table 4.** Friends versus strangers: HbO WTC in actual dyads (23 versus 23).

| Stimulus | Friends M (Md) | Strangers M (Md) | Delta<br>[95% CI] | r | p | p<sub>FDR</sub> |
|---|---|---|---|---|---|---|
| Audio | .7035 (.6990) | .6980 (.7003) | +.0056<br>[-.0060, .0172] | .09 | .598 | .742 |
| Comic | .7129 (.7165) | .7035 (.7075) | +.0094<br>[.0004, .0185] | .41 | .019 | .056 |
| Horror | .7034 (.7081) | .7068 (.7094) | -.0034<br>[-.0157, .0087] | -.06 | .742 | .742 |
| **All conditions** | **.7066 (.7080)** | **.7028 (.7013)** | **+.0038**<br>**[-.0020, .0096]** | **.23** | **.180** | **—** |

*Note.* M = mean, Md = median; Delta = Friends - Strangers; r = rank-biserial effect size from two-sided Mann-Whitney tests. Pooled row is outside the FDR family. Delta values may differ by .0001 from subtraction of the displayed rounded means owing to rounding. The displayed 95% bootstrap CIs target mean differences and are not multiple-comparison-adjusted; the Mann–Whitney tests compare ranks. The pooled row is descriptive.

**Figure 5.**
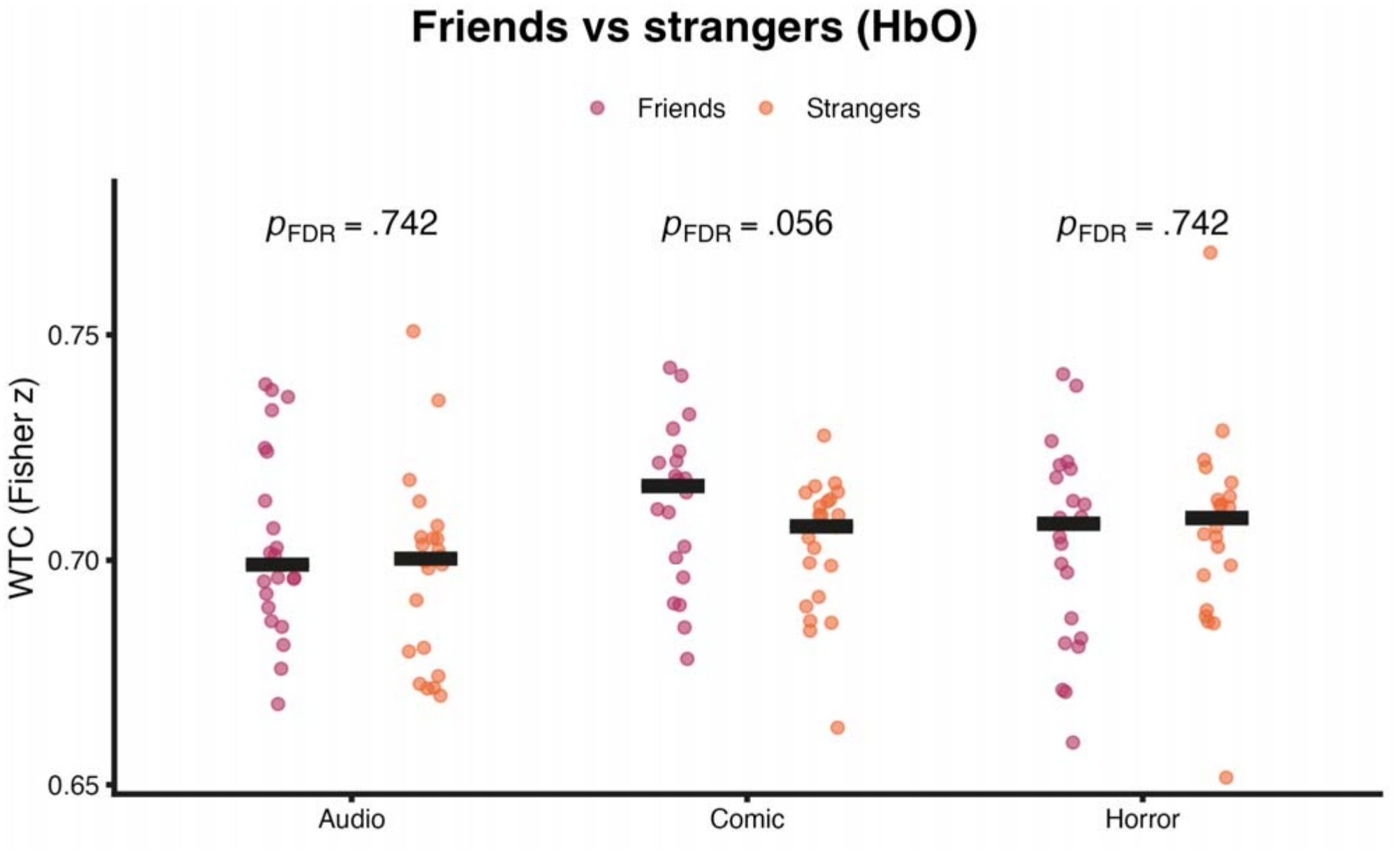
Dyad-level HbO WTC for friends and strangers (23 dyads per group). Points show individual dyads; bars indicate group medians. FDR-adjusted Mann–Whitney p values are shown above each condition. Unadjusted 95% bootstrap CIs for mean group differences are reported in Table 4; the bars in this panel represent medians, not confidence intervals.

The exploratory 60-second boundary exclusion reduced the *comic* friendship contrast from .00938 to .00688; it was no longer nominally significant (uncorrected p = .292; Supplementary Table S5). Channel and time-resolved tests also yielded no FDR-corrected familiarity effects (Supplementary Results).

HbR also showed no corrected friend-versus-stranger WTC effect (all three-stimulus FDR-adjusted p ≥ .272), including in motion sensitivity analyses (Supplementary Table S13).

### Q4 Phase structure of real-dyad coherence

Real-dyad phase angles clustered near zero in *comic* and *horror* (Rayleigh FDR-adjusted p = .0006 and .0001, respectively), but not *audio* (p = .241; Table 5 and Figure 6). Such alignment is consistent with responses to shared events and cannot identify directional influence between partners.

**Table 5.** Circular HbO phase statistics for real dyads.

| **Stimulus** | **Mean angle**<br>**[95% CI]** | **R-bar**<br>**[95% CI]** | **Rayleigh p<sub>FDR</sub>** | **V-test to 0° p<sub>FDR</sub>** |
|---|---|---|---|---|
| Audio | -18.6°<br>[-103.5°, 66.1°] | 0.18<br>[.046, .394] | .241 | .054 |
| Comic | +2.6°<br>[-21.6°, 30.4°] | 0.40<br>[.210, .607] | .0006 | .0001 |
| Horror | -1.0°<br>[-25.2°, 22.6°] | 0.46<br>[.291, .631] | .0001 | < .0001 |

*Note.* Rayleigh tests evaluate nonuniformity; the V-test evaluates orientation toward zero phase. These phase analyses were not repeated after motion control. Bracketed 95% CIs were derived by circular dyad-cluster bootstrap. Because the Audio Rayleigh test was nonsignificant, its mean-angle direction and CI should be interpreted cautiously.

**Figure 6.**
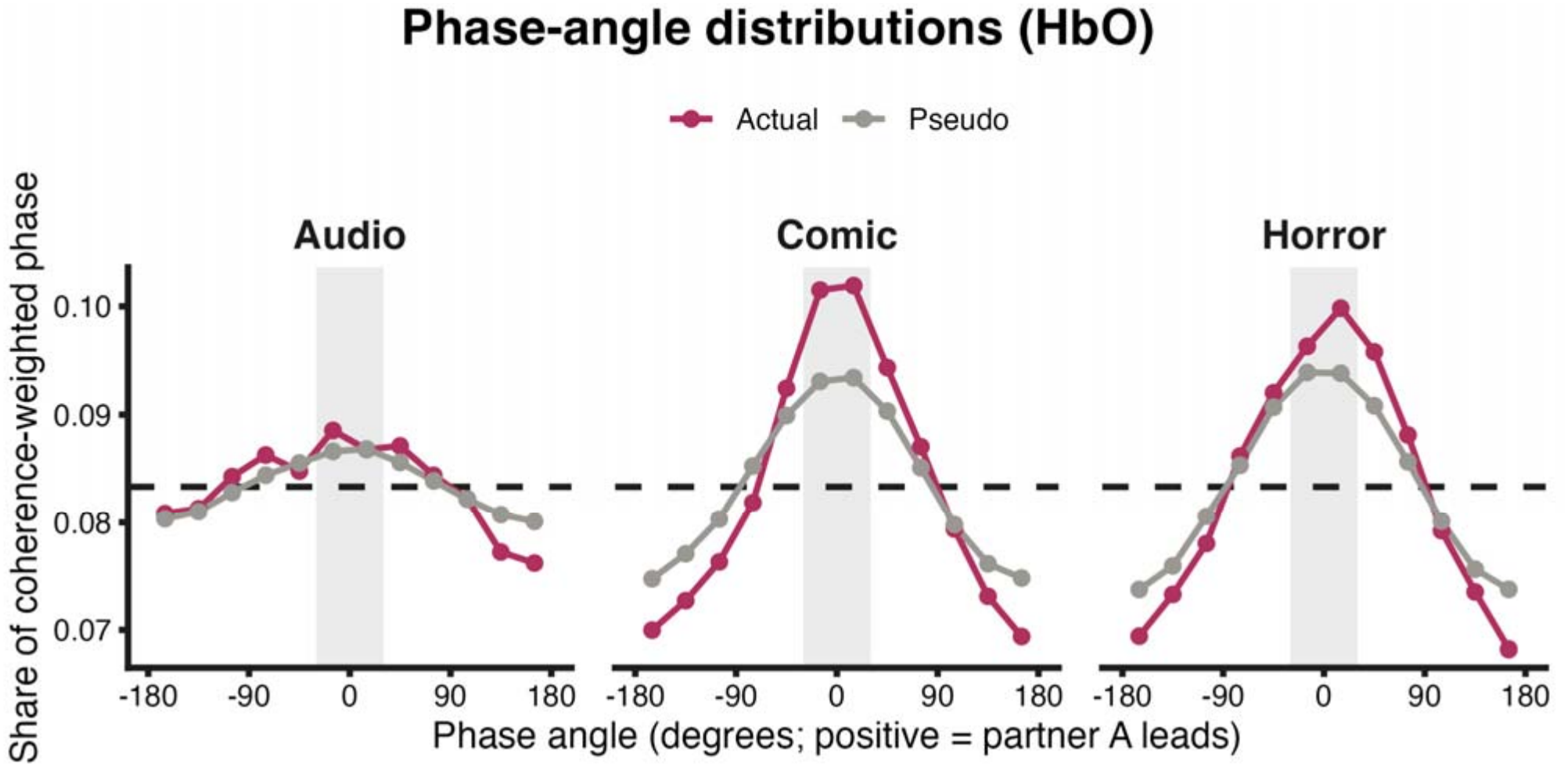
HbO phase-angle distributions in 12 bins for actual and pseudo-dyads. Curves are normalized to sum to one. Shading marks the 60° range around zero; the dashed line indicates a uniform distribution. These descriptive curves do not replace the dyad-level tests in Table 5.

Exploratory in-phase, anti-phase, and familiarity-related phase contrasts did not survive their respective FDR corrections (Supplementary Results).

In complementary HbR analyses, the *comic* V-test favored zero phase (FDR-adjusted p = .009), whereas Rayleigh nonuniformity did not survive correction (p = .056). Neither the real-versus-pseudo nor familiarity-related exploratory phase contrasts survived their 15-test corrections (Supplementary Tables S14-S15).

### Q4.1 Sensitivity to measured head motion

In 44 dyads with usable motion recordings, partners showed greater movement coherence than pseudo-dyads (all re-matching p < .001), while non-partner HbO ISC remained positive after regression of each participant’s movement measures (Supplementary Tables S6-S8).

In this subset, the *comic* real-minus-pseudo-WTC difference decreased from +.0066 (95% CI [.0022, .0109]; FDR-adjusted p = .027) before additional motion regression to +.0053 (95% CI [.0005, .0098]; adjusted p = .075) after own-motion regression. Adjustment for dyadic movement coupling gave +.0057 (95% CI [.0011, .0100]; adjusted p = .064). The bootstrap intervals and corrected re-matching tests quantify different forms of uncertainty; the results do not establish movement-independent partner-specific coherence (Supplementary Tables S7–S8).

The HbR *comic* advantage remained after own-motion regression (difference = +.0053, 95% CI [.0020, .0088]; FDR-adjusted p = .0114) and adjustment for measured between-partner motion coupling (+.0078, 95% CI [.0043, .0113]; adjusted p = .0009). Both also survived a six-test HbO/HbR correction (adjusted p = .023 and .0018, respectively). These secondary results do not exclude unmeasured movement-linked or systemic effects (Supplementary Tables S11-S12; Figure S4).

## 4 Discussion

Here, we examined cerebral hemodynamics in friend dyads and pseudo-dyads during naturalistic viewing of three different short movie types (*comic, horror, audio*). We found a small *comic*-specific increase in inter-brain hemodynamic synchrony in actual dyads relative to pseudo-dyads. The primary HbO effect survived checks of stimulus-boundary handling, but was less robust to motion adjustment. Complementary HbR results showed a similar *comic* advantage persisting after tested motion adjustments. Neither chromophore showed a consistent familiarity effect. Together, these findings support a cautious interpretation of interpersonal synchrony during co-viewing (discussed below).

Positive non-partner ISC is consistent with previous work showing similar responses to shared films and narratives (Hasson et al., 2004; Nummenmaa et al., 2012; Nastase et al., 2019). Its persistence after excluding block boundaries suggests it was not restricted to stimulus transitions. Even so, the present design cannot isolate cortical stimulus responses from shared arousal or physiology.

Everyone viewed the material with a partner, so the non-partner comparisons do not provide an independent-viewing baseline.

Actual partners showed slightly greater HbO WTC than pseudo-dyads during *comic*, even after modest boundary exclusion. Complementary HbR effect points in the same direction, remaining after regression of the recorded head-motion features. Nonetheless, HbR was analyzed retrospectively in the same participants, not in an independent replication. Both chromophores are susceptible to systemic physiology, and people recorded together share an environment and behavior. The partial reduction after motion regression and the absence of short-separation measurements leaves open whether movement artifacts, movement-linked physiology, or coordinated behavior contributed to the effect (Hamilton, 2021; Leahy et al., 2025; Yücel et al., 2021). Boundary and motion controls were not jointly applied, so these results do not isolate a neural effect specific to the partner.

We found no consistent friendship effect in either chromophore. Although the HbO *comic* friend-versus-stranger comparison was nominally positive, it did not survive FDR correction and became weaker after boundary exclusion; HbR familiarity contrasts were also nonsignificant after correction. With 23 dyads per group (22 per group with usable motion recordings), sensitivity to small effects was limited. The wide confidence intervals and corrected tests do not establish an overall familiarity effect, but nonsignificant contrasts do not demonstrate equivalence. Similar responses among friends may reflect shared interests or interpretations rather than direct interaction (Parkinson et al., 2018). Laughter is a contagious social emotion (Scott et al., 2014), and the augmented response found in the *comic* condition may reflect interpersonal processes as well.

HbO phase angles clustered near zero in *comic* and *horror*, while HbR showed a weaker *comic*-specific orientation toward zero in the V-test (but not the corrected Rayleigh test). Neither chromophore showed robust exploratory real-versus-pseudo or familiarity-related phase contrasts after correction. Phase alignment may follow shared stimulus timing or physiology; these analyses were not repeated after motion regression and cannot establish direction of neural influence.

The study lacked short-separation or cardiovascular measurements. HbO and HbR were derived from the same recordings, so cross-chromophore agreement does not represent independent replication or remove shared systemic influences. All participants were female and each stimulus category was represented by one example, limiting generalization to broader populations or genres. The primary pseudo-dyad control matched stimulus content but not every aspect of the recording session. These features limit the interpretation of the small *comic* difference. Another limitation is that our fNIRS recordings were limited to lateral frontal and frontopolar regions, and recordings extending posteriorly and laterally to parietal and temporal regions may disclose aspects of inter-regional neural synchronization within and between individuals possibly differentially engaged by this task (e.g., De Felice et al., 2024).

Future experiments should compare co-viewing with individual viewing of the same stimuli, include concurrent physiological and motion measurements, and sample multiple stimulus examples. Larger and more diverse dyadic samples would also help clarify whether familiarity has a small or context-dependent effect. Such designs could distinguish shared stimulus responses from additional coherence associated with being together.

## 5 Conclusion

This study demonstrates consistent inter-subject alignment in frontal hemodynamic responses during naturalistic co-viewing, as well as inter-brain synchrony between actual partners during the *comic* condition. The primary HbO effect was robust to the tested stimulus-boundary choices but sensitive to head-motion adjustment; complementary HbR analyses showed the same directional advantage, including after adjustment for measured movement. Neither chromophore established a consistent friendship or phase-specific effect. By combining non-partner ISC, stimulus-matched pseudo-dyads, and sensitivity analyses, these results distinguish shared stimulus responses from additional partner/session-related coherence without establishing interaction-specific neural coupling. Independent-viewing controls and concurrent physiological measurements are needed to identify the mechanisms involved. Finally, this methodology can be adapted for use with other populations, (e.g., neurodivergent ones), where shared dyadic interactions between caregivers and patients are of interest, but difficult to measure directly using self-report questionaries or third-party observational techniques.

## Supporting information

Supplementary Methods, Results, Figures and Tables

## Data and Code Availability Statement

The summary results supporting the findings of this study are presented in the article and its Supplementary Material. The de-identified fNIRS data supporting the findings of this study are publicly available on <u>Zenodo</u>. The Python code used for data preprocessing, inter-subject correlation (ISC), wavelet transform coherence (WTC), statistical analyses, and sensitivity analyses is publicly available on <u>GitHub</u>. The code repository includes documentation and software dependencies to facilitate the reproducibility of the reported analyses.

## Ethics Statement

The study involving human participants was reviewed and approved by the Research Ethics Committee of the School of Psychology at Trinity College Dublin (REAMs No. 5291; approval letter dated 19 January 2026). The study was conducted in accordance with applicable institutional and national ethical requirements. All participants provided written informed consent prior to participation.

## Author Contributions

XF: Conceptualization, Investigation, Data curation, Formal analysis, Visualization, Software, Writing – original draft, Writing – review & editing.

MB: Formal analysis, Visualization, Software, Validation, Writing – review & editing. KA: Funding acquisition, Writing – review & editing.

DT: Funding acquisition, Writing – review & editing.

SOM: Conceptualization, Supervision, Funding acquisition, Writing – review & editing.

All authors read and approved the final version of the manuscript and agree to be accountable for the work.

## Funding

This research received support through the Trinity Research Doctorate Award (lead PI, Daniela Tropea; TRDA MultiPI Award 2025 (DT)). The fNIRS equipment used in this study was purchased with funding from the Higher Education Authority Research Boost Programme (2025).

## Acknowledgments

The authors sincerely thank Delphine Guichard and Komal Zade from the Rett Research Group, Trinity College, Dublin, for their helpful suggestions. We also gratefully acknowledge the support of the Trinity Research Doctorate Award (TRDA, 2025) and the Higher Education Authority Research Boost Programme (2025), which supported the purchase of the fNIRS equipment used in this research. ChatGPT (OpenAI, GPT-6) was used to assist with manuscript language editing and formatting during the initial manuscript drafting; Claude (Anthropic, Opus 5.5) was used to assist with code writing. The authors are solely responsible for the scientific and final manuscript content.

## Conflict of Interest

The authors declare that the research here was conducted in the absence of any commercial or financial relationships that could be construed as a potential conflict of interest.

## Notes

### Competing Interest Statement

The authors have declared no competing interest.

