## Supplementary Methods, Results, Figures and Tables for "Frontal Cortical Hemodynamic Responses and Inter-Brain Synchrony During Naturalistic Co-Viewing of Differing Movie Types"

Supplementary Methods and Results

Stimulus-boundary, motion-control, and chromophore sensitivity of hemodynamic synchrony during naturalistic co-viewing

### Supplementary Methods

The boundary sensitivity analysis used 92 SNIRF recordings from 46 dyads. Each recording contained three stimulus blocks marked by alternating high (H, onset) and low (L, offset) event markers; the markers do not identify the stimulus, so each block was labelled from the recording's presentation-order code. Both partners in every dyad carried the same code and identical block onsets. The six codes were: 1 = comic, audio, horror; 2 = comic, horror, audio; 3 = horror, comic, audio; 4 = audio, comic, horror; 5 = horror, audio, comic; 6 = audio, horror, comic (eight dyads per code, except six dyads for code 4). Preprocessing followed the optical-density, SCI, TDDR, modified Beer–Lambert, and Hamming-window FIR steps described in the main Methods. Aligned stimulus segments contained 14,975 samples (approximately 600 seconds). ISC excluded actual partners; WTC was masked within the cone of influence. Central-interval analyses retained 50-second WTC windows only when fully contained after removing 30, 60, or 120 seconds from each block boundary. These overlapping post hoc thresholds were analyzed separately, without correction across thresholds.

For each condition, 10,000 one-to-one re-matchings of the 46 A and 46 B participants, excluding actual partner matches, formed the WTC null distribution. The displayed unrestricted pseudo mean represents all 2,070 non-partner A–B pairs and is descriptive, not an independent sample of 2,070 dyads. An additional sensitivity analysis allowed re-matching only within the same presentation order. This changes the null distribution and its expected mean. FDR correction was applied across the three stimuli within each trimming threshold; results across thresholds reuse the same participants.

To test whether FIR filtering depended on neighboring blocks, pre-filter HbO values inside Comic were held fixed while samples immediately outside the block, within the approximately 165-second one-sided filter support, were replaced by reflected or constant extensions. We then reapplied the same FIR filter and recalculated WTC across all 2,116 A–B combinations. This was a sensitivity check, not an alternative recommended preprocessing pipeline.

Boundary WTC was computed from HbO data sampled at approximately 5 Hz using 64 complex Morlet scales, 50-second windows, one-second steps, and cone-of-influence masking. Agreement with 6,348 full-rate 25-Hz condition-by-pair scores was close (mean absolute difference = .000032), although the sampling rates differed. Motion-regressed WTC was not recalculated after boundary exclusion.

#### Additional motion-control procedures

Both participants had usable accelerometer and gyroscope data in 44 dyads (88 people); one friend dyad and one stranger dyad lacked usable inertial recordings. Motion predictors were compared with each participant's HbO signals, with a non-partner motion signal serving as a reference for explained variance.

Sensitivity analyses compared unadjusted HbO, HbO after regression of the participant's own motion predictors, and coherence adjusted for measured between-partner motion coupling. Partner movement coherence was compared with pseudo-dyads by participant re-matching. Actual-versus-pseudo WTC used the one-to-one re-matching null; familiarity analyses used dyad-level Mann–Whitney tests. FDR correction was applied across the three stimuli within each specification. Pooled summaries were descriptive. The analyses were performed on the same participants and were not independent replications or jointly repeated with boundary trimming.

Participant sign-flip test for ISC. Participant-mean ISC values are not independent, because each pairwise ISC contributes to the means of both participants in the pair, so the Wilcoxon test can be anticonservative. The sign-flip test respects this dependence. For each draw, every participant received a random sign (+1 or −1), and each pairwise Fisher-z value was multiplied by the product of the two participants' signs, so that all pairs sharing a participant were flipped together. The test statistic was the sum of all eligible pairwise values (actual partners and channels failing SCI excluded), equivalent to their mean. Under the null hypothesis of zero mean ISC, the signs are exchangeable at the participant level. The two-sided p value was (1 + number of draws with an absolute statistic at least as large as the observed one) / (10,000 + 1), using 10,000 draws (seed [2026, 1]); the smallest attainable value is therefore approximately .0001. The same procedure was applied to each channel, to HbR, to the trimmed blocks, and to ISC after motion regression (Chen et al., 2016).

Uncertainty estimation. For analyses with participant- or dyad-level source data, two-sided 95% percentile CIs were obtained from 20,000 nonparametric bootstrap draws of intact dyads (seed [2026, 2]), stratified by familiarity (23 friends and 23 strangers; 22 per group in the IMU subset). Every real-versus-pseudo draw recomputed the actual-pair mean and the eligible between-original-dyad pseudo-pair mean; duplicate bootstrap copies of one original pair were not treated as non-partner dyads. Motion-coupling regression slopes were refitted within each draw. CIs for familiarity comparisons refer to mean differences, whereas Mann–Whitney tests target rank-based contrasts; unadjusted bootstrap CIs are not equivalent to FDR-adjusted inferences. Conservative 98.33% Bonferroni CIs were additionally checked for three-film families. Circular phase CIs used dyad-level resampling. Cluster-bootstrap CIs were not computed for the trimmed ISC and trimmed real-versus-pseudo WTC estimates.

#### Complementary HbR analysis

After inspection of the primary HbO findings, HbR was examined as a secondary, exploratory measure in the same 92 SNIRF recordings. The same SCI ≥ .70 channel criterion, TDDR, modified Beer–Lambert conversion, 0.02–0.10 Hz continuous FIR filtering, film alignment, and actual-versus-pseudo pairing rules were used. Non-partner HbR ISC used the 25-Hz series. HbR WTC used the same 64 Morlet scales, cone-of-influence mask, 50-second windows and one-second steps, at 5 Hz. All HbR phase measures, for real dyads and for the full real/pseudo comparisons, were computed at 5 Hz.

For HbR, ISC was tested by participant-level Wilcoxon and sign-flip checks; actual-versus-pseudo WTC by 10,000 partner-excluding one-to-one re-matchings; familiarity by dyad-level Mann–Whitney tests; and phase structure by Rayleigh and zero-directed V-tests. FDR correction was applied across the three stimuli for each specified Q1–Q3 comparison, and across 15 contrasts for each exploratory real/pseudo or familiarity-related Q4 phase family. A six-test family combining HbO and HbR condition-level Q2 comparisons was also examined within each motion specification. Two-sided percentile 95% CIs used 20,000 stratified intact-dyad bootstrap draws and were not multiplicity adjusted. Motion analyses in 44 dyads used the same IMU predictors and between-partner motion-coherence covariate as for HbO. HbR phase was not reanalyzed after motion regression, and HbR was not jointly recomputed with the HbO boundary-trimming analysis.

### Supplementary Results

#### Non-partner ISC after boundary trimming

**Table S1.** Non-partner ISC after excluding stimulus-boundary intervals.

| **Stimulus** | **Trim each side (s)** | **Mean ISC (z)** | **Positive** | **Wilcoxon p** | **Sign-flip p** |
| --- | --- | --- | --- | --- | --- |
| Audio | 0 | 0.01271 | 78/92 | 1e-11 | 0.0001 |
| Audio | 30 | 0.01289 | 76/92 | 3.2e-11 | 0.0001 |
| Audio | 60 | 0.01220 | 74/92 | 4.3e-10 | 0.0001 |
| Audio | 120 | 0.01229 | 73/92 | 4.3e-09 | 0.0001 |
| Comic | 0 | 0.03109 | 87/92 | 9.5e-16 | 0.0001 |
| Comic | 30 | 0.03241 | 88/92 | 8.9e-16 | 0.0001 |
| Comic | 60 | 0.03516 | 88/92 | 1e-15 | 0.0001 |
| Comic | 120 | 0.03941 | 85/92 | 3.1e-15 | 0.0001 |
| Horror | 0 | 0.04418 | 84/92 | 1.1e-15 | 0.0001 |
| Horror | 30 | 0.04695 | 85/92 | 1.6e-15 | 0.0001 |
| Horror | 60 | 0.04701 | 85/92 | 2.4e-15 | 0.0001 |
| Horror | 120 | 0.04884 | 82/92 | 7e-15 | 0.0001 |

All condition-specific Wilcoxon tests remained significant after FDR correction at every trimming threshold. The ordering of condition means is descriptive; these analyses did not test genre effects (Figure S1). The trimmed condition means are point estimates; corresponding dyad-cluster bootstrap CIs were not computed.

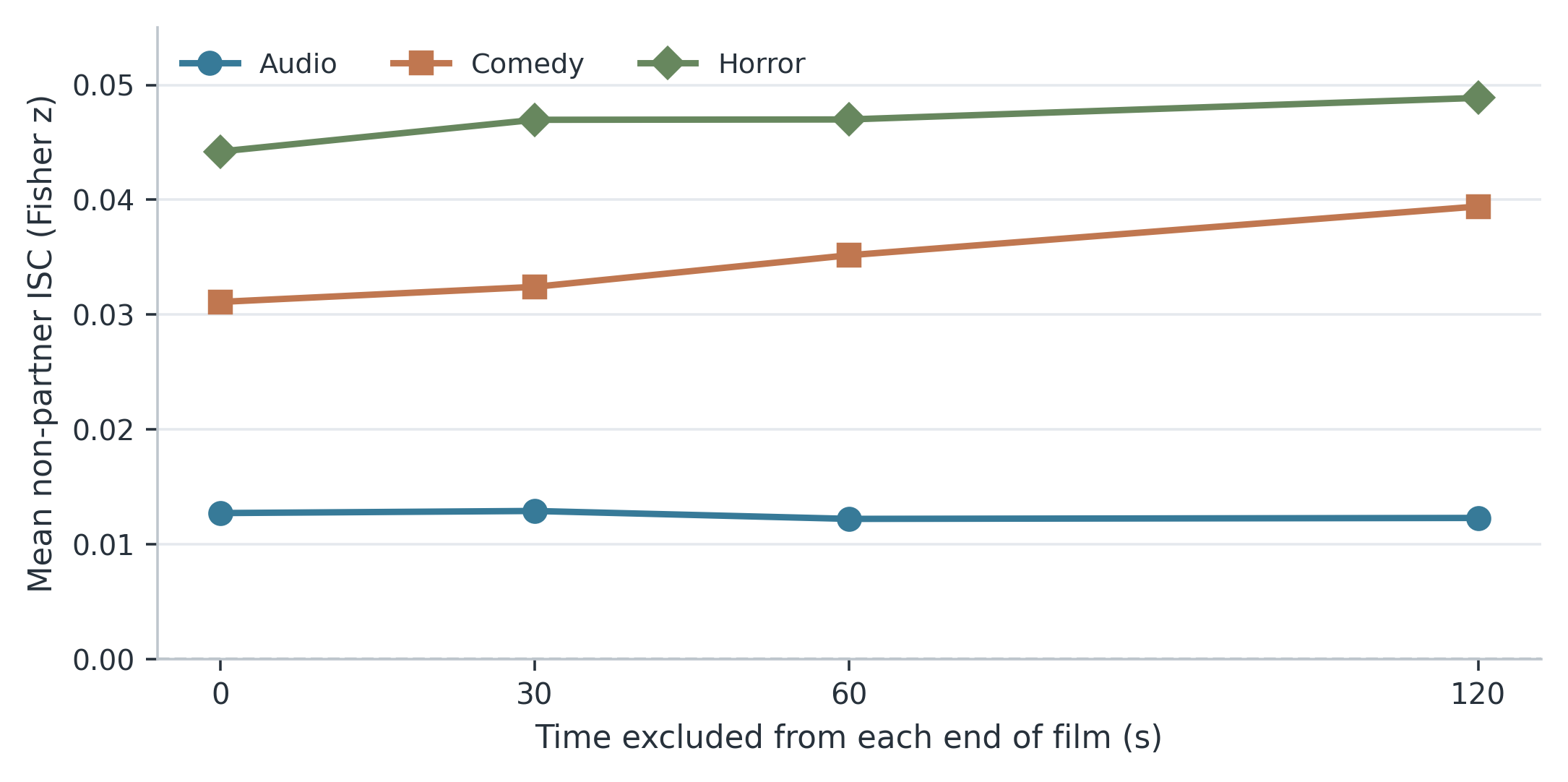

**Figure S1.** Mean non-partner ISC (Fisher z) after excluding 0, 30, 60, or 120 seconds from each stimulus boundary (n = 92). Estimates reuse participants and are not independent replications; all Wilcoxon tests remained significant after FDR correction. Curves show mean estimates only, without confidence limits.

#### Unrestricted participant re-matching (primary null)

**Table S2.** Absolute WTC in real dyads and the unrestricted non-partner pseudo pool after central-interval exclusion.

| **Stimulus** | **Trim (s)** | **Real** | **Pseudo** | **Difference** | **One-sided p** | **p_FDR_** |
| --- | --- | --- | --- | --- | --- | --- |
| Audio | 0 | 0.70072 | 0.69897 | +0.00175 | 0.2454 | 0.2454 |
| Audio | 30 | 0.69678 | 0.69588 | +0.00090 | 0.3653 | 0.3653 |
| Audio | 60 | 0.69443 | 0.69399 | +0.00044 | 0.4305 | 0.4305 |
| Audio | 120 | 0.69331 | 0.69385 | -0.00054 | 0.5555 | 0.5555 |
| Comic | 0 | 0.70821 | 0.70207 | +0.00615 | 0.0120 | 0.0360 |
| Comic | 30 | 0.70645 | 0.69992 | +0.00653 | 0.0110 | 0.0330 |
| Comic | 60 | 0.70564 | 0.69885 | +0.00679 | 0.0122 | 0.0366 |
| Comic | 120 | 0.70503 | 0.69810 | +0.00693 | 0.0248 | 0.0744 |
| Horror | 0 | 0.70509 | 0.70073 | +0.00437 | 0.0554 | 0.0831 |
| Horror | 30 | 0.70323 | 0.69856 | +0.00467 | 0.0515 | 0.0772 |
| Horror | 60 | 0.70162 | 0.69669 | +0.00494 | 0.0490 | 0.0735 |
| Horror | 120 | 0.70056 | 0.69713 | +0.00343 | 0.1592 | 0.2388 |

#### Stimulus-order-matched re-matching (exploratory null)

**Table S3.** Exploratory re-matching null restricted to the same stimulus presentation order.

| **Stimulus** | **Trim (s)** | **Real** | **Null mean** | **Real - null** | **One-sided p** | **p_FDR_** |
| --- | --- | --- | --- | --- | --- | --- |
| Audio | 0 | 0.70072 | 0.69730 | +0.00342 | 0.0731 | 0.0731 |
| Audio | 30 | 0.69678 | 0.69418 | +0.00260 | 0.1460 | 0.1460 |
| Audio | 60 | 0.69443 | 0.69217 | +0.00226 | 0.1993 | 0.1993 |
| Audio | 120 | 0.69331 | 0.69233 | +0.00098 | 0.3757 | 0.3757 |
| Comic | 0 | 0.70821 | 0.70222 | +0.00599 | 0.0073 | 0.0219 |
| Comic | 30 | 0.70645 | 0.70005 | +0.00640 | 0.0057 | 0.0171 |
| Comic | 60 | 0.70564 | 0.69894 | +0.00670 | 0.0071 | 0.0213 |
| Comic | 120 | 0.70503 | 0.69770 | +0.00732 | 0.0119 | 0.0357 |
| Horror | 0 | 0.70509 | 0.70048 | +0.00461 | 0.0379 | 0.0568 |
| Horror | 30 | 0.70323 | 0.69823 | +0.00500 | 0.0301 | 0.0451 |
| Horror | 60 | 0.70162 | 0.69642 | +0.00521 | 0.0312 | 0.0468 |
| Horror | 120 | 0.70056 | 0.69683 | +0.00373 | 0.1162 | 0.1743 |

Order restriction changes the permutation null. The differences in Table S3 therefore should not be compared directly with the unrestricted pseudo mean in Table S2; the analyses use the same dyads. Dyad-cluster CIs were not computed for Tables S2–S4.

#### Comic: sensitivity to exterior FIR input

**Table S4.** Comic FIR exterior-input sensitivity checks (46 × 46 A–B combinations).

| **Exterior treatment** | **Trim each side (s)** | **Real** | **Pseudo** | **Difference** | **Uncorrected p** |
| --- | --- | --- | --- | --- | --- |
| Continuous filtering | 0 | 0.70821 | 0.70207 | +0.00615 | 0.0120 |
| Continuous filtering | 60 | 0.70564 | 0.69885 | +0.00679 | 0.0122 |
| Reflected exterior | 0 | 0.70770 | 0.70151 | +0.00619 | 0.0114 |
| Reflected exterior | 60 | 0.70566 | 0.69884 | +0.00682 | 0.0122 |
| Constant exterior | 0 | 0.70907 | 0.70275 | +0.00632 | 0.0102 |
| Constant exterior | 60 | 0.70593 | 0.69910 | +0.00683 | 0.0121 |

Each exterior-input treatment used 10,000 re-matchings, excluding actual partners. Reflected and constant extensions were tested only for Comic; their p values were not adjusted across three stimuli.

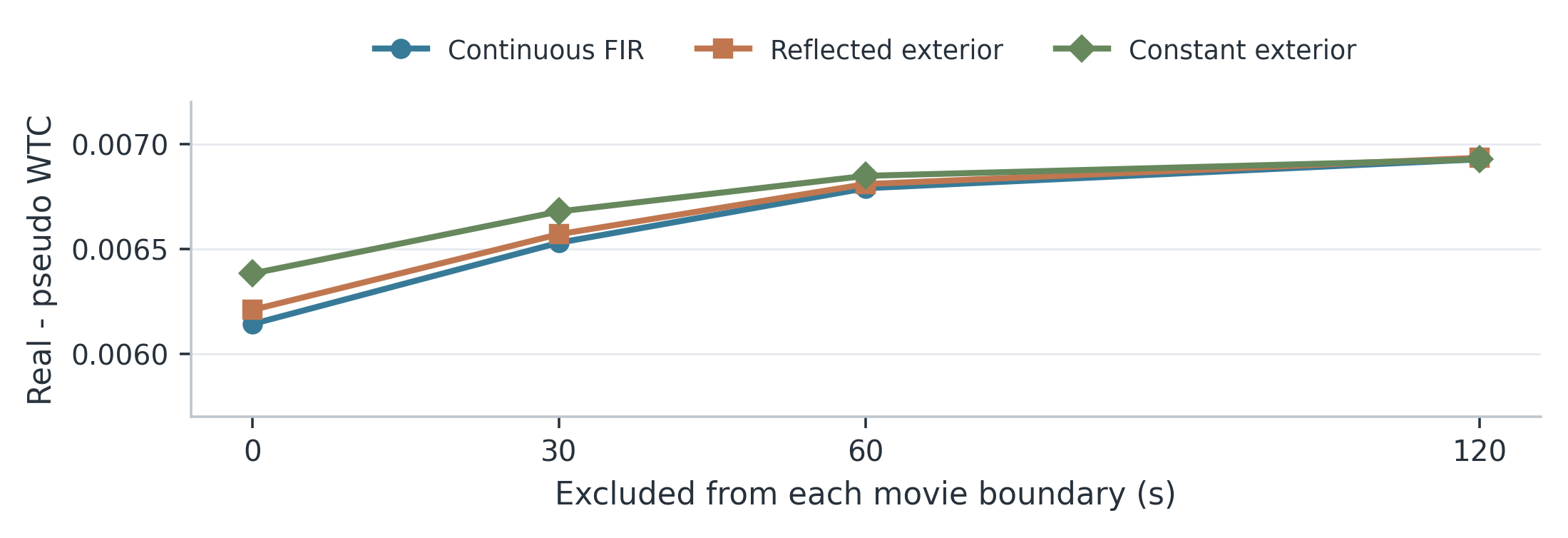

**Figure S2.** Comic actual-minus-pseudo WTC after boundary trimming under continuous FIR filtering and reflected or constant exterior input. Points are descriptive estimates, not confidence intervals. Table S4 gives numerical values for the 0- and 60-second thresholds.

#### Friends versus strangers after stimulus-boundary exclusion

**Table S5.** Friends-versus-strangers WTC differences among actual dyads (23 versus 23) after exclusion of boundary intervals.

| **Stimulus** | **Trim each side (s)** | **Friends - strangers [95% CI]** | **Two-sided p** | **p_FDR_** |
| --- | --- | --- | --- | --- |
| Audio | 0 | +0.00557 [-.0060, .0171] | 0.5828 | 0.7252 |
| Audio | 30 | +0.00420 [-.0073, .0157] | 0.7088 | 0.9125 |
| Audio | 60 | +0.00482 [-.0068, .0166] | 0.7088 | 0.9475 |
| Audio | 120 | +0.00291 [-.0100, .0158] | 0.7252 | 0.7921 |
| Comic | 0 | +0.00937 [.0006, .0184] | 0.0187 | 0.0562 |
| Comic | 30 | +0.00797 [-.0020, .0178] | 0.0866 | 0.2598 |
| Comic | 60 | +0.00688 [-.0039, .0181] | 0.2916 | 0.8749 |
| Comic | 120 | +0.00798 [-.0065, .0224] | 0.3448 | 0.7921 |
| Horror | 0 | -0.00340 [-.0156, .0088] | 0.7252 | 0.7252 |
| Horror | 30 | -0.00345 [-.0165, .0093] | 0.9125 | 0.9125 |
| Horror | 60 | -0.00217 [-.0155, .0109] | 0.9475 | 0.9475 |
| Horror | 120 | +0.00035 [-.0134, .0139] | 0.7921 | 0.7921 |

The boundary results do not establish a stable friendship effect. Dyad-level Mann–Whitney tests used one WTC summary per actual pair; trimming thresholds overlap and were not adjusted as a separate family. Motion-regressed WTC was not recalculated after boundary removal. The 5-Hz boundary WTC estimates closely matched, but were not identical to, the 25-Hz scores. Bracketed 95% percentile bootstrap CIs are for friend-minus-stranger mean WTC, not adjusted for repeated trim thresholds or the three stimuli; Mann–Whitney p values remain rank based.

#### Exploratory channel, time, and phase comparisons

Among familiarity tests, four of 72 condition-by-channel comparisons were nominally significant, but none survived FDR correction (minimum adjusted p = .121). Of 1,650 sliding-window tests, 31 had uncorrected p < .05, and none survived correction (minimum adjusted p = .92). The real-versus-pseudo in-phase contrast in Comic (.300 versus .278; uncorrected p = .003) did not survive correction (adjusted p = .051); neither did the anti-phase contrast (uncorrected p = .044; adjusted p = .27). During Audio, friends showed nominally higher in-phase (r = .43, p = .014) and lower anti-phase coherence (r = −.43, p = .012) than strangers, but neither contrast survived FDR correction (both adjusted p = .10). Other familiarity-related phase contrasts had p ≥ .22. These exploratory comparisons involved the same participants as the main analyses.

#### Motion coupling and motion-control sensitivity analyses

##### Motion-related variation in HbO and interpersonal movement

Own head motion explained 13.6%, 15.2%, and 14.7% of HbO variance during Audio, Comic, and Horror, compared with 8.8%, 8.9%, and 9.6% for another participant's motion signal (t(87) = 11.4, 12.0, and 11.7, respectively; Table S6). Movement coherence was greater for actual partners than pseudo-dyads (re-matching z = 6.7, 4.5, and 6.9; all p < .001). Movement energy also showed cross-participant ISC (z = .046, .065, and .125; all p < 10⁻¹⁴).

**Table S6.** Variance in HbO associated with head motion (n = 88 participants).

| **Stimulus** | **Own motion** | **Other-person motion** | **t(87)** |
| --- | --- | --- | --- |
| Audio | 13.6% | 8.8% | 11.4 |
| Comic | 15.2% | 8.9% | 12.0 |
| Horror | 14.7% | 9.6% | 11.7 |

Note. Explained variance is from motion models; another participant's motion served as a reference. The excess variance explained by own versus another participant's motion was 4.9 percentage points for Audio (95% CI [4.0, 5.7]), 6.3 for Comic ([5.3, 7.3]), and 5.1 for Horror ([4.2, 6.0]); dyad-cluster bootstrap.

Across paired recordings, movement coherence was weakly associated with HbO coherence in Comic (r = .08) and Horror (r = .11; corrected p = .005). This does not establish causal motion contamination. Mean movement magnitude did not differ reliably between friends and strangers (all Mann–Whitney p ≥ .23) and was not associated with dyad WTC (all p ≥ .40).

##### Persistence of ISC after own-motion regression

Non-partner ISC remained positive after regression of each participant's motion features (n = 88): mean Fisher z = .0125 for Audio (95% CI [.0073, .0183]; Wilcoxon p = 3.4 × 10⁻¹¹), .0291 for Comic (95% CI [.0211, .0380]; p = 2.4 × 10⁻¹⁵), and .0346 for Horror (95% CI [.0250, .0454]; p = 9.7 × 10⁻¹⁵). Sign-flip p = .0001 in all conditions. Corresponding estimates before regression were .0123, .0308, and .0430. Motion regression attenuated Horror ISC most strongly but did not remove the positive cross-participant associations.

##### Real-versus-pseudo and familiarity contrasts after movement adjustment

The Comic real-versus-pseudo difference was +.0066 before additional motion adjustment (FDR-adjusted p = .027), +.0053 after own-motion regression (adjusted p = .075), and +.0057 after adjusting for between-partner motion coupling (adjusted p = .064; Table S7). The pooled difference declined from +.0040 (p = .007) to +.0013 (p = .200) after own-motion regression; movement-coupling adjustment gave +.0028 (p = .037). Pooled values were not included in the three-stimulus FDR family. Movement-independent partner coherence was therefore not established. Despite lower CI limits above zero for the Comic mean WTC differences, the original motion-adjusted re-matching tests did not pass three-stimulus FDR correction; these results should not be called independently significant.

Boundary and motion sensitivity analyses addressed separate questions and were not combined. Greater stability under boundary trimming does not establish independence from movement.

**Table S7.** Motion sensitivity of real-minus-pseudo absolute WTC (44 dyads).

| **Stimulus** | **HbO unchanged: Delta (p)** | **Own motion removed: Delta (p)** | **Motion coupling adjusted: Delta (p)** |
| --- | --- | --- | --- |
| Audio | +.0012 (.321) [-.0049, .0076] | -.0026 (.861) [-.0077, .0024] | +.0006 (.412) [-.0054, .0066] |
| Comic | +.0066 (.009) [.0022, .0109] | +.0053 (.025) [.0005, .0098] | +.0057 (.021) [.0011, .0100] |
| Horror | +.0042 (.065) [-.0019, .0103] | +.0012 (.323) [-.0043, .0064] | +.0022 (.209) [-.0034, .0078] |
| **All conditions** | **+.0040 (.007) [.0008, .0072]** | **+.0013 (.200) [-.0011, .0036]** | **+.0028 (.037) [-.0002, .0057]** |

Note. Numbers in parentheses are uncorrected one-tailed p-values. Comic p_FDR_ = .027 (unchanged), .075 (own motion removed), and .064 (motion coupling adjusted). Pooled values are outside the correction families. Bracketed values are unadjusted two-sided 95% dyad-cluster bootstrap CIs for mean differences (20,000 resamples); these do not invert the one-sided re-matching tests.

Familiarity contrasts also varied with the motion-control specification. In the 44-dyad subset after own-motion regression, friends had higher WTC during Comic (+.0105; rank-biserial r = .38; FDR-adjusted p = .048), whereas strangers had higher WTC during Horror (−.0114; r = −.40; FDR-adjusted p = .048). Neither difference survived correction in unchanged HbO or after adjustment for dyadic motion coupling. The direction and significance of the familiarity comparison therefore depended on the analysis specification.

**Table S8.** Motion sensitivity of friend-minus-stranger absolute WTC (44 dyads).

| **Stimulus** | **HbO unchanged: Delta (p)** | **Own motion removed: Delta (p)** | **Motion coupling adjusted: Delta (p)** |
| --- | --- | --- | --- |
| Audio | +.0055 (.647) [-.0065, .0177] | +.0066 (.201) [-.0035, .0167] | +.0056 (.716) [-.0065, .0174] |
| Comic | +.0083 (.042) [-.0009, .0175] | +.0105 (.032) [-.00004, .0207] | +.0081 (.045) [-.0010, .0171] |
| Horror | -.0043 (.565) [-.0169, .0082] | -.0114 (.024) [-.0230, .0002] | -.0047 (.716) [-.0171, .0066] |
| **All conditions** | **+.0032 (.286) [-.0028, .0091]** | **+.0019 (.549) [-.0029, .0068]** | **+.0030 (.255) [-.0026, .0084]** |

Note. Numbers in parentheses are uncorrected two-sided Mann–Whitney p-values. Only the own-motion-removed Comic and Horror contrasts pass their within-family FDR (p_FDR_ = .048 each); other versions do not. The pooled row is outside these families. Bracketed values are unadjusted 95% bootstrap CIs for mean differences; the p values are from rank-based tests. In Comic and Horror after own-motion regression the mean-difference CIs extend slightly across zero despite the corrected rank-test p values.

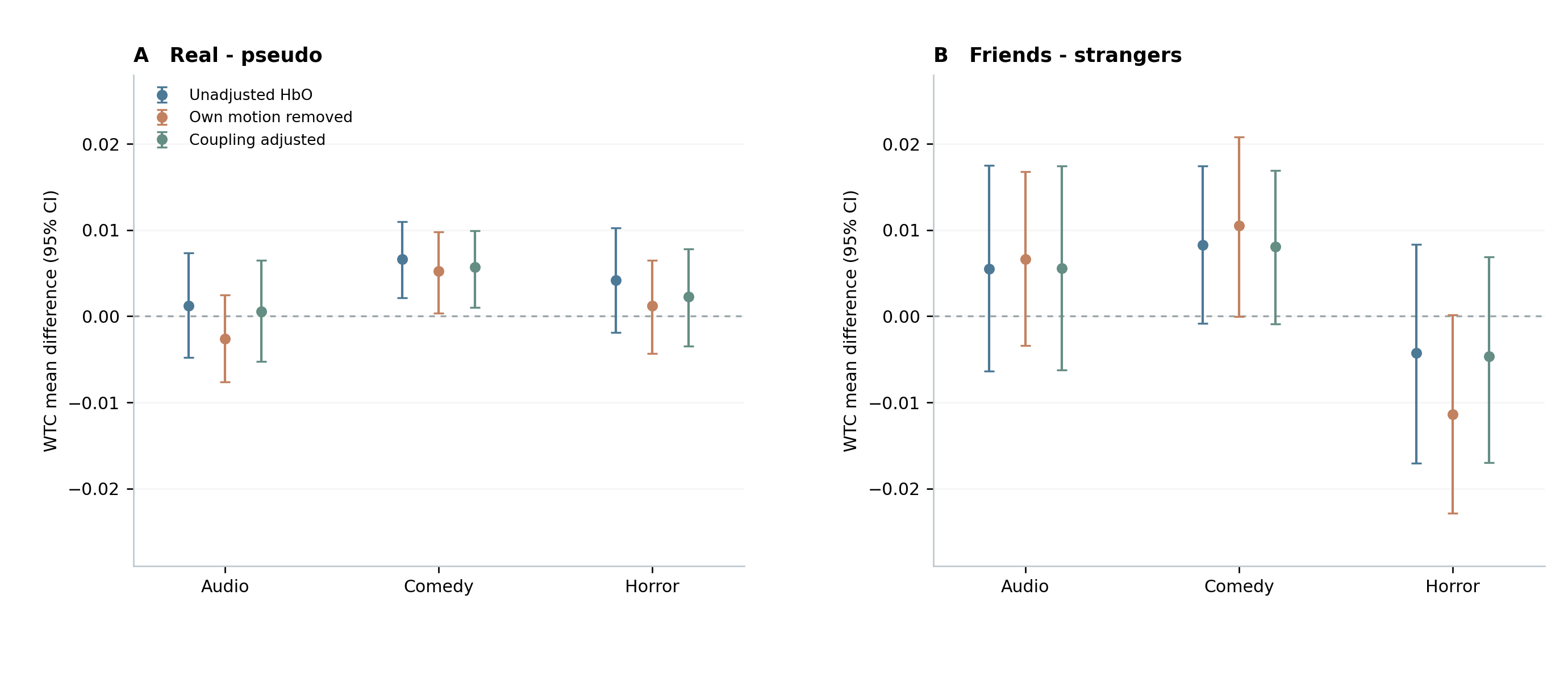

**Figure S3.** Point estimates and 95% dyad-cluster bootstrap CIs for (A) real-minus-pseudo WTC and (B) friend-minus-stranger mean WTC by stimulus and motion-control specification (44 dyads). Intervals are unadjusted; original inferential tests and corrected p values are reported in Supplementary Tables S7 and S8.

#### Complementary HbR analyses (secondary and exploratory)

HbR was derived from the same recordings as HbO and is not an independent replication. The analyses below use the same participants and pseudo-dyad construction as the primary results; all displayed intervals are unadjusted 95% CIs.

##### Non-partner HbR ISC (Q1)

Non-partner HbR ISC remained positive in Audio, Comic, and Horror. Each condition passed the three-stimulus FDR correction, and sign-flip p = .0001 throughout (Table S9). Of 72 condition-by-channel comparisons, 63 passed the Wilcoxon FDR and 61 passed the sign-flip FDR.

**Table S9.** Non-partner HbR ISC across all 92 participants.

| **Stimulus** | **Mean Fisher z [95% CI]** | **Positive** | **Wilcoxon pFDR** | **Sign-flip p** |
| --- | --- | --- | --- | --- |
| Audio | .0054 [.0020, .0091] | 69/92 | 5.7e-09 | .0001 |
| Comic | .0105 [.0067, .0145] | 80/92 | 7.4e-13 | .0001 |
| Horror | .0214 [.0145, .0291] | 85/92 | 7.4e-13 | .0001 |

Note. The 95% intervals use intact-dyad bootstrap resampling; Wilcoxon and sign-flip tests assess participant-level positive ISC and are not derived by inverting these intervals.

##### HbR actual-versus-pseudo WTC (Q2)

An HbR actual-partner coherence advantage was detected only in Comic (Table S10). The Comic result remained below .05 after a six-test FDR correction across HbO and HbR condition-level Q2 comparisons. Audio and Horror did not survive their three-stimulus correction.

**Table S10.** Real-minus-pseudo HbR WTC (46 dyads; 2,070 descriptive pseudo-pairs).

| **Stimulus** | **Real** | **Pseudo** | **Delta [95% CI]** | **p** | **pFDR (3)** | **pFDR (6)** |
| --- | --- | --- | --- | --- | --- | --- |
| Audio | .6988 | .6979 | +.0009 [-.0028, .0046] | .3097 | .3097 | .3097 |
| Comic | .7076 | .6989 | +.0088 [.0054, .0123] | .0001 | .0003 | .0006 |
| Horror | .7024 | .6998 | +.0026 [-.0015, .0067] | .1113 | .1669 | .1669 |

Note. p is the one-sided, one-to-one re-matching probability. pFDR (3) is corrected across HbR stimuli; pFDR (6) includes the three corresponding HbO comparisons. Pseudo-pairings were not treated as independent observations. The HbR WTC values are computed at 5 Hz.

##### HbR motion-control analyses

HbR ISC remained positive after regression of each participant’s recorded motion features. The Comic HbR real-minus-pseudo contrast was reduced by own-motion regression but remained significant after correction within the tested motion specifications (Table S11). Motion models explained more HbR variance when using each participant’s own movement than when using an unrelated participant’s movement (Table S12). These covariates do not remove unmeasured systemic physiology.

**Table S11.** HbR real-minus-pseudo WTC across motion-control specifications (44 dyads).

| **Stimulus** | **Motion specification** | **Delta [95% CI]** | **pFDR (3)** | **pFDR (6)** |
| --- | --- | --- | --- | --- |
| Audio | Unchanged | +.0009 [-.0029, .0048] | .3095 | .3221 |
| Audio | Own motion regressed | -.0011 [-.0051, .0027] | .7277 | .8532 |
| Audio | Motion coupling adjusted | -.0001 [-.0039, .0036] | .5261 | .5261 |
| Comic | Unchanged | +.0085 [.0050, .0121] | .0003 | .0006 |
| Comic | Own motion regressed | +.0053 [.0020, .0088] | .0114 | .0228 |
| Comic | Motion coupling adjusted | +.0078 [.0043, .0113] | .0009 | .0018 |
| Horror | Unchanged | +.0023 [-.0020, .0066] | .2058 | .2058 |
| Horror | Own motion regressed | +.0035 [-.0008, .0077] | .0675 | .0900 |
| Horror | Motion coupling adjusted | +.0005 [-.0036, .0046] | .5261 | .4981 |

Note. All p values originate from one-sided partner-excluding re-matching tests. Each model has a separate three-stimulus FDR family; six-test corrections include HbO and HbR within the same model. CIs are unadjusted dyad-cluster bootstrap intervals. The 44-dyad subset differs from the full 46-dyad analysis.

**Table S12.** Head-motion associations and post-regression HbR ISC (88 participants).

| **Stimulus** | **Own motion HbR R²** | **Unrelated motion HbR R²** | **Own - unrelated R² (pp) [95% CI]** | **Cleaned ISC z [95% CI]** |
| --- | --- | --- | --- | --- |
| Audio | 12.9% | 9.0% | 3.95 [3.35, 4.57] | .0042 [.0012, .0074] |
| Comic | 14.0% | 9.4% | 4.60 [3.90, 5.30] | .0089 [.0051, .0132] |
| Horror | 13.4% | 9.7% | 3.67 [2.83, 4.54] | .0179 [.0118, .0247] |

Note. pp = percentage points. Paired comparisons of explained HbR variance had p < .0001 in each film. Post-regression HbR ISC also remained positive (Wilcoxon FDR-adjusted p < .0001; participant sign-flip p = .0001 for each film). These are associations with the measured IMU covariates, not proof of causal artifact removal.

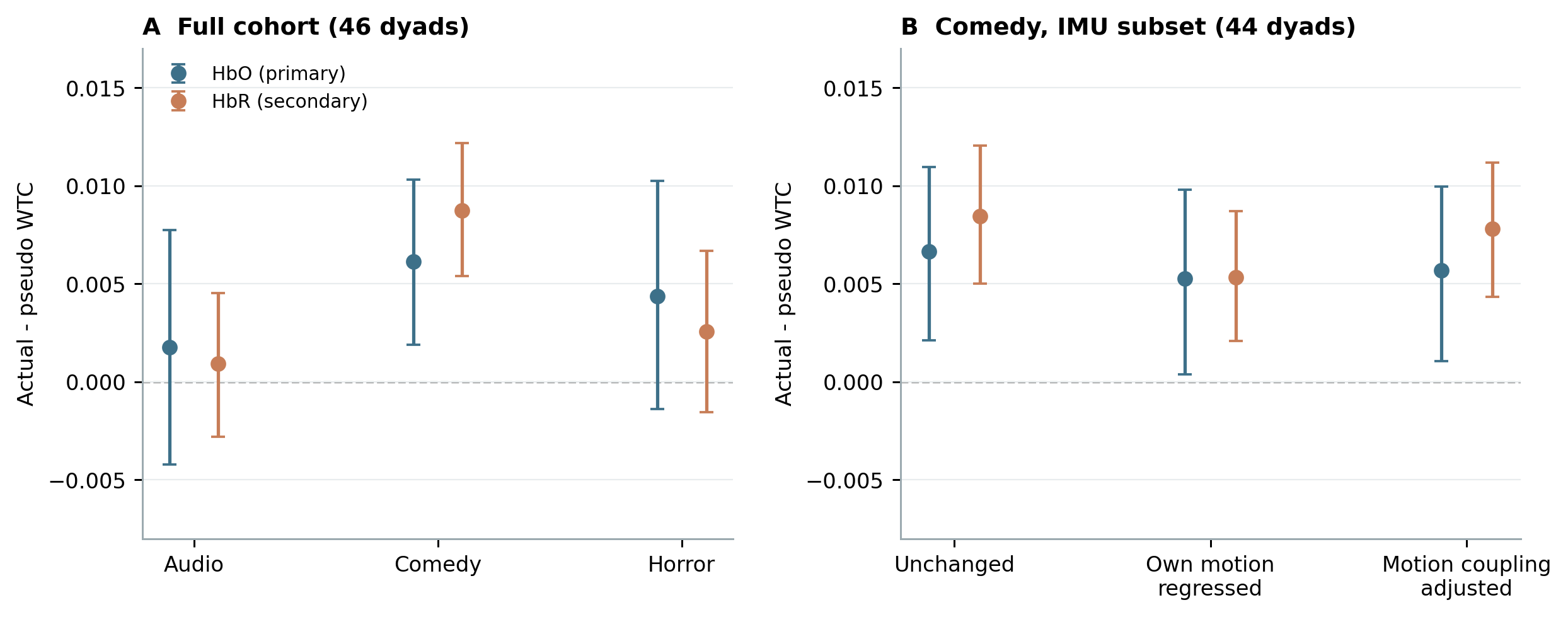

**Figure S4.** Real-minus-pseudo WTC effects for HbO (primary) and HbR (secondary). (A) Full-cohort estimates by stimulus (46 dyads). (B) Comic by motion-control specification in the 44-dyad IMU subset. Points and error bars show effect means and unadjusted 95% dyad-cluster bootstrap CIs; corrected re-matching p values are in main-text Table 2 and Supplementary Tables S7, S10, and S11. The chromophores come from the same recordings.

##### HbR social familiarity (Q3)

No friend-versus-stranger HbR WTC difference survived three-stimulus FDR correction (Table S13). None of 72 channel comparisons or 1,650 overlapping-window comparisons survived their respective FDR corrections. Likewise, none of the HbR familiarity contrasts passed FDR correction in the 44-dyad movement subset under any of the three motion-control specifications (minimum adjusted p = .366).

**Table S13.** HbR WTC in friends and strangers (23 dyads per group).

| **Stimulus** | **Friends M** | **Strangers M** | **Delta [95% CI]** | **p** | **pFDR** |
| --- | --- | --- | --- | --- | --- |
| Audio | .7005 | .6970 | +.0035 [-.0041, .0111] | .2817 | .2817 |
| Comic | .7110 | .7043 | +.0067 [-.0010, .0144] | .0907 | .2722 |
| Horror | .6986 | .7061 | -.0075 [-.0167, .0013] | .2355 | .2817 |

Note. Delta = mean friends - strangers HbR WTC. The p values are from two-sided Mann–Whitney rank tests; the bootstrap CIs target mean differences and are not multiplicity-adjusted. Cross-chromophore consistency does not establish equivalence between groups.

##### HbR phase organization (Q4)

Among actual dyads, the zero-directed V-test detected a Comic phase orientation (FDR-adjusted p = .0092), but the Comic Rayleigh test did not survive correction (p = .0560; Table S14). Thus, the phase evidence is weaker than the Comic WTC-magnitude contrast. No real-versus-pseudo phase measure passed correction across 15 comparisons (Table S15), nor did the 15 exploratory familiarity-phase contrasts (minimum adjusted p = .195). HbR phase analyses were not repeated after motion regression.

**Table S14.** Circular phase statistics for actual HbR dyads (n = 46).

| **Stimulus** | **Mean angle [95% CI]** | **R-bar [95% CI]** | **Rayleigh pFDR** | **V-test to 0° pFDR** |
| --- | --- | --- | --- | --- |
| Audio | -32.4° [-198.6°, +134.5°] | 0.030 [.023, .282] | .9602 | .4044 |
| Comic | +12.4° [-30.4°, +54.0°] | 0.293 [.128, .491] | .0560 | .0092 |
| Horror | -21.8° [-81.1°, +52.8°] | 0.193 [.064, .392] | .2723 | .0645 |

Note. The V-test targets the specified zero-phase direction; Rayleigh tests general nonuniformity. The unwrapped Audio mean-angle CI crosses the circular branch cut and is not informative as a conventional linear interval because R-bar is near zero.

**Table S15.** Exploratory HbR real-minus-pseudo phase contrasts (15-test FDR family).

| **Stimulus** | **Phase measure** | **Real - pseudo [95% CI]** | **Two-sided p** | **pFDR (15)** |
| --- | --- | --- | --- | --- |
| Audio | In-phase proportion | -.0040 [-.0127, .0046] | .4182 | .6016 |
| Audio | Anti-phase proportion | +.0053 [-.0031, .0137] | .2690 | .5739 |
| Audio | Lagged proportion | -.0013 [-.0087, .0059] | .7127 | .8258 |
| Audio | Phase consistency | -.0092 [-.0199, .0018] | .1578 | .4734 |
| Audio | Mean lag (s) | +.004 [-.206, .212] | .9759 | .9899 |
| Comic | In-phase proportion | +.0078 [-.0040, .0195] | .1440 | .4734 |
| Comic | Anti-phase proportion | -.0107 [-.0219, .0006] | .0284 | .2130 |
| Comic | Lagged proportion | +.0029 [-.0051, .0107] | .4412 | .6016 |
| Comic | Phase consistency | +.0180 [.0028, .0333] | .0082 | .1230 |
| Comic | Mean lag (s) | +.163 [-.044, .380] | .1034 | .4734 |
| Horror | In-phase proportion | -.0053 [-.0151, .0049] | .3324 | .5739 |
| Horror | Anti-phase proportion | +.0017 [-.0075, .0110] | .7157 | .8258 |
| Horror | Lagged proportion | +.0035 [-.0045, .0113] | .3444 | .5739 |
| Horror | Phase consistency | -.0084 [-.0226, .0069] | .2544 | .5739 |
| Horror | Mean lag (s) | +.002 [-.204, .207] | .9899 | .9899 |

Note. Re-matching tests are two-sided and exclude actual partners. The 15-test FDR family includes five measures in each of the three stimuli. The intervals are unadjusted 20,000-draw dyad-cluster bootstrap CIs; some may exclude zero despite nonsignificant FDR-adjusted permutation tests.
